# Diversity, abundance, and biogeography of CO_2_ fixing microorganisms in peatlands

**DOI:** 10.1101/2025.02.07.637021

**Authors:** Marie Le Geay, Kyle Mayers, Anna Sytiuk, Ellen Dorrepaal, Martin Küttim, Mariusz Lamentowicz, Eeva-Stiina Tuittila, Béatrice Lauga, Vincent E.J. Jassey

## Abstract

Microbial communities play a crucial role in the carbon (C) dynamic of peatlands— a major terrestrial C reservoir. While heterotrophic microorganisms attracted much attention over the past decades due to their role in peatland greenhouse gases emissions, CO_2_ fixing microorganisms (CFMs) remained particularly overlooked. Here, by leveraging metabarcoding and digital droplet PCR (ddPCR), we provide a comprehensive survey of CFM communities, including oxygenic phototrophs, chemoautotrophs and aerobic anoxygenic phototrophic bacteria (AAnPBs), in different peatland types. We demonstrate that CFMs are very abundant and diverse in peatlands, with on average 1021 CFMs contributing up to 40% of the total bacterial abundance. In particular, we show that oxygenic phototrophs (mostly Cyanophyceae and Palmophylloceae) are the most abundant CFMs, closely followed by chemoautotrophs (Proteobacteria) and AAnPBs (Vulcanimicrobiia). Using a joint-species distribution model, we further find that CFMs aggregate into six major clusters with different niche size. These clusters constitute the core and specific CFM microbiome. The core microbiome, which the occurrence is strongly influenced by temperature and nutrients, directly modulate the diversity and abundance of CFMs. Our findings highlight the importance of CFM diversity and abundance in peatlands, further reveal their complex structuration in link with environmental conditions and suggest that changes in environmental conditions could shift CFMs communities. These results are the foundation to better understand the role of CFMs for the peatland C cycle inputs.

## Introduction

Peatlands are a massive terrestrial carbon (C) sink, storing about 30% of all soil C (500 - 1,000 Gt of C) for only 3% of land area (Nichols and Peteet, 2019; Yu et al., 2021). This storage of C hinges on the imbalance between C loss through respiration and decomposition and C uptake through photosynthesis. Cold temperature, waterlogged condition and poor nutrient availability (Rydin and Jeglum, 2006) preserve organic matter from decomposition, making peatlands an essential component for mitigating climate changes (Strack et al., 2022). Microbial growth and activity are important players of the peatland C dynamics as microorganisms are involved in key processes of the peatland C cycle such as decomposition, respiration, methanogenesis and CO_2_ fixation (Dorrepaal et al., 2009; Hopple et al., 2020; Hamard et al., 2021b; Juottonen et al., 2022). Because climate warming is expected to accelerate the microbially driven efflux of CO_2_ toward the atmosphere (Davidson and Janssens, 2006; Bardgett et al., 2008), heterotrophic microorganisms received increasing attention over the last decades (Dorrepaal et al., 2009; Bell et al., 2018; Gavazov et al., 2018; Hopple et al., 2020; Gao et al., 2022) whilst CO_2_ fixing microorganisms (CFMs) remained overlooked. Yet, CFM could balance, to some extent, microbial CO_2_ emissions in response to warming (Le Geay et al., 2024a).

CFMs can fix atmospheric CO_2_ through seven natural metabolic pathways including the Calvin-Benson-Bassham (CBB) cycle (reductive pentose phosphate cycle), the rTCA cycle (reductive citrate cycle), the 3-HP/4-HB cycle (3-hydroxypropionate/4-hydroxybutyrate cycle ), the 3-HP cycle (3-hydroxypropionate bi-cycle), the Wood–Ljungdahl pathway (the reductive acetyl-CoA pathway), the DC/4-HB cycle (dicarboxylate hydroxybutyrate cycle) and the reductive glycine pathway (Berg, 2011; Bar-Even et al., 2012; Figueroa et al., 2018; Sánchez-Andrea et al., 2020; Huang et al., 2022). Among these metabolic pathways, the CBB cycle was the first discovered and is the most widespread (Bassham et al., 1954; Falkowski et al., 2008). In particular, the CBB cycle is found in all oxygenic phototrophs and in chemoautotrophs (Yuan et al., 2012; Bay et al., 2021). Oxygenic phototrophs fix CO_2_ by harnessing the energy of light, through photosynthesis while chemoautotrophs power the CBB cycle by oxidizing chemicals or molecular hydrogen (Hügler and Sievert, 2011; Gomez-Saez et al., 2017; Wang et al., 2023). Recent work further showed that some aerobic anoxygenic phototrophic bacteria (AAnPBs) possess and express CBB genes, suggesting they fix atmospheric CO_2_ too (Graham et al., 2018; Yabe et al., 2022). While the mechanisms of microbial CO_2_ fixation through the CBB cycle are well documented, the main microbial diversity and abundance associated with CFMs using the CBB cycle remain poorly known in soils, and especially in peatlands.

Oxygenic phototrophs can be abundant and diverse in peatlands (Gilbert and Mitchell, 2006; Hamard et al., 2021b), living either in the pore water, in the water film on *Sphagnum* mosses or in the hyaline *Sphagnum* cells that store water (Gilbert et al., 1998). They are sensitive to temperature, soil water content and light availability as well as plant cover, dissolved organic carbon and pH (Hamard et al., 2021b, 2021a; Richy et al., 2024). On the contrary, chemoautotrophs and AAnPBs have been overlooked while they can be as abundant as oxygenic phototrophs (Le Geay et al., 2024b). As oxygenic phototrophs are essential for the peatland C cycle (Hamard et al., 2021a) and because chemoautotrophs and AAnPBs could play an important role too (Gios et al., 2024; Le Geay et al., 2024b), better understanding the diversity, community composition, abundance and sensitivity to environmental parameters of these CFMs is essential to improve our comprehension of the peatland C cycle.

In this study, we combined a metabarcoding approach with digital droplet PCR (ddPCR) to explore how the abundance, diversity and community structure of CFM using the CBB cycle vary across peatland types and depth. In particular, we aim to identify which environmental variables drive CFMs attributes and their interplay. Because of the strong variations in abiotic factors with depth and between peatland types, we hypothesize that (i) patterns of CFM diversity and community composition will vary between peatlands sites and that (ii) such changes are mainly driven by nutrients, acidity and temperature along with precipitation as these parameters are strong drivers of microbial communities (Yuan et al., 2012; Guo et al., 2015; Nowak et al., 2015; Lew et al., 2016). Given their sensitivity to light availability and oxygen content, we also hypothesize that (iii) oxygenic phototrophs and AAnPBs will prevail in the upper peat layers whereas chemoautotrophs will be favored in deeper layers where reduced compound are more abundant (Rydin and Jeglum, 2006; Frei et al., 2012; Bai et al., 2015; Hädrich et al., 2019). Finally, following the recent findings of Le Geay et al., 2024b showing the simultaneous presence of oxygenic phototrophs, chemoautotrophs and AAnPBs in peat samples, we propose that (iv) these functional groups will contribute to the peatland C fixation together instead of only oxygenic phototrophs.

## Materials and methods

### Study sites and sampling

We selected four peatlands along a latitudinal gradient spanning different environmental conditions and trophic states, from northern Sweden to southern France. Counozouls (42°41’16N - 2°14’18E; 1.350 m above sea level (a.s.l)) is a moderately rich fen in the Pyrenees mountains, southwestern France. This site is characterized by a mean annual temperature (MAT) and annual precipitation (MAP) of 7.1°C and 1,027 mm, respectively. Männikjärve (58°52’30N - 26°15’04E; 82 m a.s.l) is an ombrotrophic bog situated in Central Estonia in the Endla mire system and characterized by MAT of 5.9°C and MAP of 623 mm. Siikaneva (61°50’00N - 24°11’21E; 160 m a.s.l) is a boreal oligotrophic fen in southern Finland with MAT and MAP of 5.1°C and 611 mm, respectively. Abisko (68°20’54N - 19°04’09E; 350 m a.s.l) is a palsa mire characterized by MAT of 2.7°C and MAP of 418 mm. More details on these sites can be found in Sytiuk et al. (2022).

At each site, we collected five peat cores (10 cm diameter, 20 cm depth) in homogeneous *Sphagnum* habitats during summer 2022. Peat cores were further subdivided into three depths (D1: 0-5 cm; D2: 5-10 cm and D3: 10-15 cm), corresponding to the living, decaying and dead layers, respectively (Le Geay et al., 2024b). At each depth, a few grams of peat were collected, cut in small species, homogenized, and placed in sterile 5 mL Eppendorf tubes containing 3 mL of RNA*later* (ThermoFisher) for environmental DNA analysis. The remaining peat was placed in sterile plastic bags for chemical analyses. We also sampled pore water to complete the chemical characterization. All samples were stored at -20°C prior proceeding to DNA extraction and chemical analyses.

### Collection of environmental data

Daily air and soil temperature, precipitation and water table depth were measured at hourly intervals at each site since 2019 (Meters® sensors and data loggers; Meter Group, Pullman, WA, USA). Average air temperature, soil temperature, precipitation and water table depth for the spring period (20^th^ of March – 21^st^ of June) and for the winter period (21^th^ of December – 19^th^ of March) were calculated using these data. Pore water was collected in piezometers at each site and filtered through 0.45 *µ*m pore size (Whatman) to measure pH, dissolved organic carbon (DOC), dissolved total nitrogen (DTN) and the quality of the dissolved organic matter (DOCq). DOC and DTN were measured by combustion on a Shimadzu TOC-L. DOCq was assessed by measuring the aromatic content and molecular weight of DOC. To do so, we measured the absorbance of DOC between 250 and 660 nm (15 wavelengths in total) in 200-µL sample aliquots in 96-well quartz microplate using a BioTek SynergyMX spectrofluorometer (Jaffrain et al., 2007). Demineralized water filtered through Whatman filter was used as a blank and to correct the values. We then retrieved RFE (relative fluorescence efficiency) , freshness, FI (fluorescence index), peak A, C, M and BIX (biological index) to assess the DOCq. Peat soil water content was measured by weighing peat samples fresh and dry (lyophilized samples; expressed per g of H_2_O per g of dry peat (gH_2_O.g.-1 DW)). Peat samples were further used to measure several cations (Li^+^, Na^+^, NH_4_^+^, K^+^, Mg^2+^ and Ca^2+^) and anions (F^-^, Cl^-^, NO_2_^-^, NO_3_^-^, Br^-^, SO_4_^2-^ and PO_4_^3-^) using high performance ion chromatography on a Dionex DX-120 and on a Dionex ICS-5000+ respectively. Finally, we quantified total concentrations of carbohydrates, flavonoids, tannins, phenols and water phenols in the peat following Sytiuk et al., (2022).

### DNA extraction, amplification and sequencing

DNA was extracted using the DNeasy PowerSoil Pro Kit (Qiagen) following manufacturer’s instructions. DNA concentration was quantified using a Nanodrop ND-1000 spectrophotometer. Extracts were then stored at -20°C prior to DNA amplification. For all samples we targeted four different genes, namely the 16S rRNA gene, the 23S rRNA gene, *cbbL* and *bchY*. The 16S rRNA gene allowed to investigate all prokaryotes while the others genes were used to target CFMs using the CBB cycle: the 23S rRNA gene for microorganisms involved in oxygenic photosynthesis, *cbbL* gene encoding the large subunit of RuBisCO form IA for chemoautotrophs (Kusian and Bowien, 1997; Alfreider and Bogensperger, 2018) and *bchY* encoding the Y subunit of bacteriochlorophyll biosynthesis for AAnPBs (Yutin et al., 2009). The primers pairs used for DNA amplification (Table S1.a) were PCR1-515F/PCR1-909R for the 16S rDNA gene (Wang and Qian, 2009), P23SrV-f1/P23Srv-1 for the 23S rDNA gene (Sherwood and Presting, 2007), cbbL-IA- CHEM/cbbL-IA-r for *cbbL* (Alfreider and Bogensperger, 2018) and bchY-fwd/bchY-rev to target *bchY* (Yutin et al., 2009). Each primers contained additional Illumina adapters and tags. PCRs were performed in a total volume of 50 μL containing 25 μL of AmpliTaq Gold^TM^ Master Mix (applied biosystem, ThermoFisher), 21 or 19 μL of ultrapure water, 1 μL of forward and 1 μL of reverse primer (final concentration of 20 μM) and 2 or 4 μL of DNA template according to initial DNA concentration. The PCR reaction conditions were different according to each primer pair (Table S2). PCR quality was assessed using 1.65% agarose gel electrophoresis. The high throughput sequencing was performed by the GeT-PlaGe plateform (Genotoul, Toulouse, France) using Illumina MiSeq technology with the V3 chemistry. To avoid contamination as much as possible, all molecular work was carried out in a dedicated laboratory using UV-decontaminated and sterile materials.

### Analysis of CFM communities

Sequences obtained after sequencing were already demultiplexed, trimmed of barcodes and Illumina adapters. The paired-end fastq sequences were analyzed within the FROGs pipeline v4.1.0 provided by the Galaxy web platform (Escudié et al., 2018). Paired-end sequences were merged using *VSEARCH* v2.17.2 (Rognes et al., 2016). Sequences were then denoised, dereplicated, clustered into ASVs using SWARM clustering method (Mahé et al., 2014) and chimera were removed. These ASVs were further filtered to keep only ASVs with a minimum prevalence of 2 and assigned at different taxonomic levels using the database *SILVA 138.1* (Quast et al., 2013) for the 16S rRNA gene and *µgreen db algae v1.2* (Djemiel et al., 2020) for the 23S rRNA gene. For *cbbL* and *bchY* no databases were available, affiliations were, therefore, done manually. First, sequences were aligned using Clustal OMEGA (EMBL), clustered using the *hclust* function (threshold = 0.05) and for each cluster a consensus sequence was built. Each consensus sequence was then blasted using nucleotide BLAST for highly similar sequences (megablast). Only assignations with high score, query cover (> 90%), percentage identity (> 90%) and low E-value (< 0.01), were kept while other sequences were considered as “*unassigned*”. All ASVs obtained were rarefied using *rarefy_even_depth* function of the *Phyloseq* package v1.44.0 to obtain comparable sequencing depth across samples (Fig.S1).

### Absolute quantification using digital droplet PCR (ddPCR)

Absolute quantification of genes targeting prokaryotes, oxygenic phototrophs, chemoautotrophs and AAnPBs was measured using digital droplet PCR (ddPCR, BioRad) as described in Le Geay et al., (2024). Primers pairs used for ddPCR (Table S1.b) were L/Prba338f with K/Prun518r (Øvreås et al., 1997) for prokaryotes, 16SCF/16SUR (Oh et al., 2012) for cyanobacteria, 23S255f/ P23SrV_r1(Sherwood and Presting, 2007; Le Geay et al., 2024b) for green algae and cyanobacteria, cbbLR1F/cbbLR1inR (Selesi et al., 2007) for chemoautotrophs, and pufMforward557/pufMreverse750 (Du et al., 2006) for AAnPBs. We did not used *bchY* primers to quantify AAnPBs because they were too long and too degenerated to be used in ddPCR (Achenbach et al., 2001; Béjà et al., 2002; Yutin et al., 2009). The ddPCR reactions were run on a DX200 instrument (BioRad) in a total volume of 20 µL with 10 µL of EvaGreen Supermix (BioRad, 1X), 0.5 µL of each primer (final concentration 25 µM) and 4 µL of ultrapure water. Template DNA was diluted either at 1/10, 1/100 or 1/1,000 and 5 µL were added to the reactional mix. We then used the QX200 Droplet Generator (BioRad) with QX200 Droplet Generation Oil for EvaGreen (BioRad) to emulsify the reaction mix and transferred this reactional mix into a 96-well PCR plate. The plate was heat-sealed with a foil seal and then placed on a C1000 Touch Thermocycler with deep-well module (BioRad) to run the PCR (detailed PCR programs are described in Table S3). Following amplification, plates were equilibrated for at least 10 minutes at room temperature. Then, the fluorescence was read on a QX200 Droplet Reader (BioRad) and the QuantaSoft software was used to set the threshold and analyze the results. Using ultrapure water as a negative control and different DNA extracted from cultures of microorganisms as positive controls we manually defined the threshold for each ddPCR run. *Escherichia coli* and *Micromonas pusilla* DNA diluted 1/100 were used as positive controls for prokaryotes and oxygenic phototrophs respectively. Calculation of the final concentration considered the volume of eluted DNA (70 µL), the volume of ddPCR reaction mixture (20 µL), the volume of template DNA (5 µL) and the dilution factor of the template DNA (10, 100 or 1,000). Results were further normalized using the amount of dry peat used for DNA extraction to obtain a final concentration in target copies.g^-1^ of dry peat (copies.g^-1^ DW).

### Statistical analysis

All statistical analyses were performed using Rstudio v12.0 with R build under v4.3.2 with packages specified below and graphical representations were done using *ggplot2* v3.5.1 and *igraph* v1.4.3.

Environmental drivers of samples distribution were tested by conducting a principal component analysis (PCA) using the *PCA* function of *FactoMineR* package v2.11 including all the environmental parameters. Environmental parameters were logged or squared transformed when necessary. For further analyses, environmental parameters went through a selection of variables to retain only the most significant and least collinear representative using forward selection with the function *ordiR2steps* (*vegan* v2.6.4) for climatic variables, nutrients, metabolites and organic matter quality variables. Collinearity was further assessed using *corrplot* of the *corrplot v0.92* package (Fig.S2). Correlation between absolute quantification of oxygenic phototrophs, chemoautotrophs and AAnPBs genes and environmental variables was assessed using *corrplot* of the *corrplot v0.92* package.

For each CFM gene marker (23S rRNA, *cbbL* and *bchY* genes), richness and alpha-diversity were estimated using Chao1 (*vegan* v2.6.4) and Shannon (*vegan* v2.6.4) metrics, respectively. We used two-way analysis of variance (ANOVA) to test the impact of peatland site and depth on these diversity metrics. Post-hoc tests (Tuckey honestly significant difference test) were used to identify the significative differences among factors and their interactions. To estimate species turnover between sites and depths, we also conducted a non-metric multidimensional scaling (NMDS) based on beta-diversity metric (Bray-Curtis dissimilatory). We further used PERMANOVA (permutational multivariate analysis of variance) to test for structural difference in the communities according to location and depth using the function *adonis2* of the *vegan* package v2.6-4. Correlation between selected environmental variables and diversity metrics was done using Pearson correlation.

Multiple factor analysis (MFA) was used to test the link between the ASVs matrices of oxygenic phototrophs, chemoautotrophs and AAnPBs. We chose to perform MFA as it allows to assess the general structures of datasets by coupling several groups of variables (Escofier and Pagès, 1994). First, ASVs were transformed using Hellinger transformation. Then, MFA was done using the *MFA* function of the package FactoMineR v2.11. Correlation between the three matrices were measured using RV coefficient (Pearson’s correlation coefficient). Euclidian distances of the global PCA were further used to perform cluster analysis using *hclust* with the Ward D2 method. The resulting dendrogram was plotted in the MFA ordination space.

To test the impact of location and depth on the relative abundance of CFMs, ASVs were aggregated by taxonomic rank and the relative abundance of each group (Class and Family) was calculated. Based on these relative abundances at each location and for each depth, heatmaps were generated using *geom_tile* of the ggplot2 package. Linear mixed effect models (LME) using the *lme* function of the *nlme* package v3.1-164 were implemented with location and depth as fixed explanatory terms to explain the relative abundance of each group and replicates were used as random term.

To assess the structuration of ASVs and their interaction with environmental parameters, we used joint species distribution modelling (JSDM) (Warton et al., 2015). First, we standardized environmental parameters using the *decostand* function of the vegan package v2.6.4 and we filtered the ASVs to keep only ASVs with a minimum of 0.1% in relative abundance. Then, we ran the JSDM model using the *JSDM_binomial_probit* function of the jSDM package v0.2.6 on presence/absence data (Warton et al., 2015). The model was checked using diagnostic plots and we got the correlation of environmental data using *get_enviro_cor* function and the residual correlation using *get_residual_cor* function of the JSDM package v0.2.6. We then built the co-occurrence network using the strongest correlation link and using the function *cluster_fast_greedy* of the igraph package. We also recovered the beta-coefficient of the JSDM model and plot these coefficients for each identified cluster. For each cluster we further analyzed the taxonomic affiliation of the ASVs belonging to the cluster. The relative abundances of the six main clusters generated by the JSDM package were recovered for each sample and projected in the MFA ordination space.

The size of niche spaces of ASVs in each cluster was estimated by calculating hypervolumes using *hypervolumes* R package (Blonder, 2018; Chen et al., 2024). We defined the n-dimensional hypervolume of each ASV using the site scores of the first three axes of the PCA describing environmental conditions in each site.

Finaly, to explore the main drivers of CFM community structure and gene abundances we built a set of path diagrams subjected to structural equation modeling (SEM; Grace et al., 2015).

Adjusted R^2^ and Akaike information criterion were used to compare the explanatory power of each model. First, we created two latent variables. Using results from the PCR and the environmental factors selection we tested several environmental parameters to design the latent variable “Environment” that represents the impact of the main environmental drivers. The best predictors were pH, dissolved organic C (DOC), K and spring soil temperature (SST). Using the results from ddPCR we then design the latent variable “CFMs abundances” representing the global abundance of CFMs. Once the latent variables defined, we first tested the direct effect of “Environment” on clusters 1, 2 and 3 relative abundance, on species richness (Chao1), on microbial community structure (first axis of MFA) and on “CFM abundances”. Second, we tested the indirect effect of “Environment” though the interaction between clusters, between Chao1 and MFA1 and between clusters with Chao1 and MFA1 and between clusters, Chao1, MFA1 and “CFM abundances”. The paths of the SEM were fitted using the function *sem* of the lavaan package v0.6-17 (Rosseel, 2012). The adequacy of the model was determined using the *X^2^* test (P > 0.05), CFI index (comparative fit index, CFI > 0.9), AIC (Akaike value, lowest), RMSEA index (root square mean error of approximation, RMSEA < 0.05) and the RMR index (root mean square residuals, RMR < 0.05 ; Jonsson and Wardle, 2009; Grace et al., 2010).

## Results

### Partitioning of peatland sites according to environmental parameters

The four European peatland sites exhibited very different environmental conditions (Fig.1; Fig.S3-S6; Table.S4). The PCA analysis revealed a net separation of peatland sites caused by environmental parameters with both axis separating samples among peatland sites and depth (Fig.1a). The first axis separated samples between sites and was mainly driven by climatic conditions including soil temperature, air temperature, precipitations and pH (Fig.1b). These environmental variables globally followed a gradient from south (Counozouls; highest values) to north (Abisko; lowest values; Table S4; Fig.S3). The second axis separating samples between sites and depths was mainly driven by nutrients (Fig.1b). Some nutrients concentrations varied between sites (PO_4_^2-^ and NO_3_^-^) while others varied with depth (K^+^, Br^-^ and F^-^; Fig.S5). Metabolites and index of organic matter quality also contributed to both axis with few variations between sites and depth (Fig.S4 and S6).

**Figure 1.**
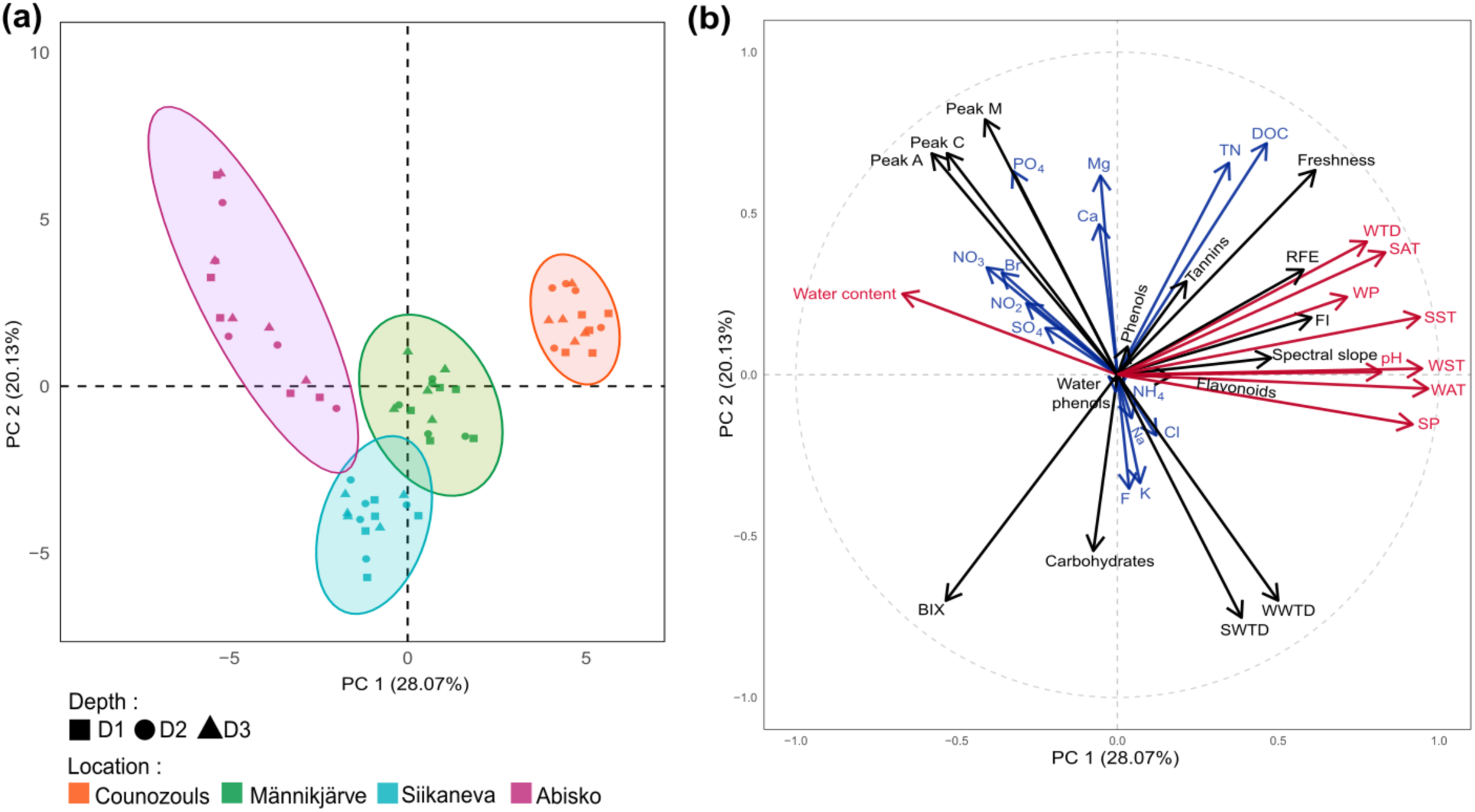
Principal component analysis (PCA) of all environmental variables with. **(a)** sample positioning in the PCA space and **(b)** environmental variables contribution to PC axis 1 and 2. Blue arrow and text represent nutrient variables and red arrow and text represent variables related to climatic condition as well as pH. DOC = dissolved organic carbon, TN = total nitrogen, RFE = relative fluorescence efficiency, BIX = biological index, FI = fluorescence index, SST = spring soil temperature, SAT = spring air temperature, SP = spring precipitation, SWTD = spring water table depth, WST = winter soil temperature, WAT = winter air temperature, WP = winter precipitation and WWTD = winter water table depth.

### Effects of depth, site and environment on CFMs abundance

CFMs were very abundant contributing up to 40% of the total bacterial abundance (Fig.2a). The 23S rRNA, *cbbL* and *pufM* genes showed similar abundances, ranging from 7.89 x 10^4^ to 3.79 x 10^6^ copies.g^-1^ of dry peat for oxygenic phototrophs, from 3.74 x 10^4^ to 3.2 x 10^6^ copies.g^-1^ of dry peat for chemoautotrophs and from 5.31 x 10^4^ to 2.44 x 10^6^ copies.g^-1^ of dry peat for AAnPBs. When compared to the 16S rRNA gene abundance we found ratios of oxygenic phototrophs, chemoautotrophs and AAnPBs very similar. Ratios of oxygenic phototrophs (16S rRNA gene/23S rRNA gene) ranged from 9% (Männikjärve, Siikaneva) to 14% (Abisko) and 20% (Counozouls) while ratios of chemoautotrophs (16S rRNA gene/*cbbL* gene) ranged from 8% (Männikjärve, Siikaneva, Abisko) to 10% (Counozouls) and ratios of AAnPBs (16Sr rRNA gene/*pufM* gene) ranged from 7% (Männikjärve, Siikaneva, Abisko) to 12% (Counozouls). Globally, we found a slight impact of location on CFM total abundance (sum of 23S rRNA, *cbbL* and *pufM* gene abundances; Fig.2b, *P* < 0.05) and at the depth level, we did not found differences (Fig.S7).

Oxygenic phototrophs abundance (23S rRNA gene) was very similar for the four sites and mostly decreased with depth (Fig.S7b). Among oxygenic phototrophs, cyanobacteria represented ∼50 % of the abundance and decreased with depth (Fig.S8). Chemoautotrophs abundance (*cbbL* gene) differed between site with higher abundance in Counozouls and Abisko and did not vary much with depth (Fig.S7c). AAnPBs abundance (*pufM* gene) was also higher in Counozouls and Abisko and decreased with depth (Fig.S7d). Environment was structuring as shown in the PCA, which allowed us to select variables impacting CFMs gene abundance as shown in Figure 2b. Oxygenic phototrophs gene abundance was notably affected by Br^-^ (*P* < 0.05; Fig.2d) while chemoautotrophs gene abundance was affected by several nutrients (PO_4_^2-^, DOC and TN) and climate parameters (pH, SST, *P* < 0.05; Fig.2d) and AAnPBs gene abundance was affected by several nutrients (PO_4_^2-^, DOC and TN) and by SST (*P* < 0.05; Fig.2d).

**Figure 2.**
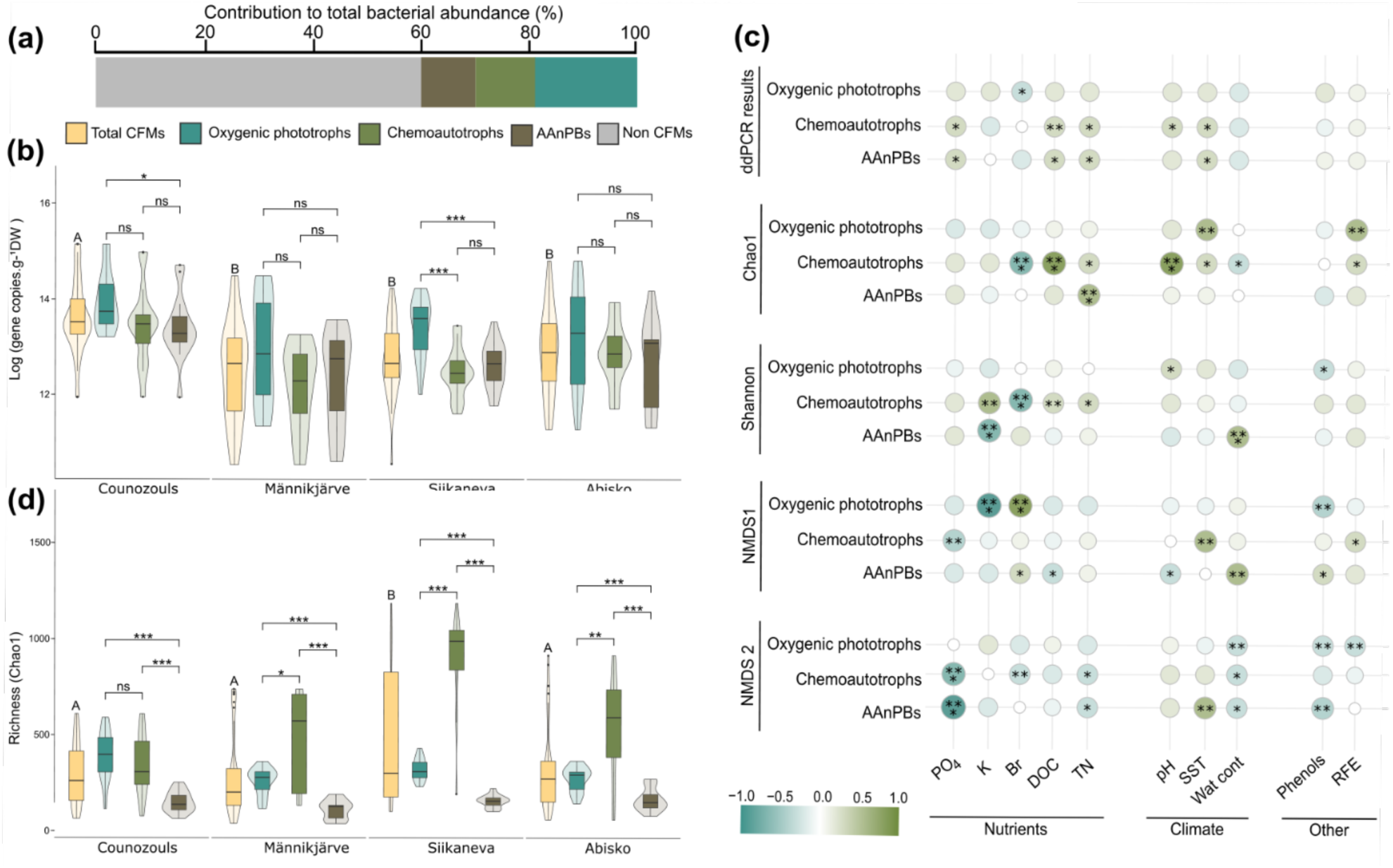
CFMs abundance, richness and their response to environmental variables. **(a)** Contribution of the different CFMs to total bacterial abundance 916S rRNA gene)**. (b)** Absolute quantification using ddPCR and **(b)** Richness (Chao1 index) of total CFMs (sum of 23S rRNA, *cbbL* and *pufM*/*bchYgenes*), oxygenic phototrophs (23S rRNA gene), chemoautotrophs (*cbbL*) and AAnPBs (*pufM/bchY*) for each site. Violin plots are showing the data distribution shape while boxplots are representing the logarithm of the total gene copies.g^-1^ DW. D1 = 0-5 cm; D2 = 5-10 cm and D3 = 10-15 cm. **(c)** Correlation plot (Pearson correlation) of CFMs community abundance (ddPCR results), richness (Chao1), alpha-diversity (Shannon) and beta-diversity (NMDS1 and NMDS2) with nutrients (PO_4_^2-^, K, Br^-^, DOC, TN), climate (pH, SST, wat cont), phenols and RFE. DOC = dissolved organic carbon; TN = total nitrogen; SST = spring soil temperature and wat cont = water content. ns = not significant; * = 0.05 P < 0.01; ** = 0.01 < P < 0.001 and *** = P < 0.001.

### Effects of depth and site on CFMs richness, diversity and community composition

In total we found 7,960 ASVs among which 2,690 belonged to oxygenic phototrophs for a mean richness of 312, 3,879 ASVs belonged to chemoautotrophs for a mean richness of 563 and the remaining 1,391 ASVs belonged to AAnPBs for a mean richness of 146. The total richness of CFMs (Chao1 index) was relatively similar among sites, at the exception of Siikaneva which was slightly higher compared to other sites (*P* < 0.05; Fig.2c). This increase was driven by a strong increase of chemoautotrophic richness in Siikaneva when compared to other sites (Fig. 2c, *P* < 0.05). CFM richness also globally decrease with depth (Fig.S9a, b and c). In term of alpha-diversity (Shannon index) we found a few variations between site (Fig.S9d, e and f). For oxygenic phototrophs we found an increase of the alpha-diversity with depth (Fig.S9d) while for chemoautotrophs and AAnPBs, alpha-diversity decreased with depth (Fig.S9e and f). The NMDS analysis further confirmed that CFMs communities were well structured both by site and depth (Fig.S10). Chemoautotrophic and aerobic anoxygenic phototrophic communities of Siikaneva and Männikjärve were notably close (Fig.S10 b and c). Richness together with alpha diversity and community structuration were very sensitive to environmental parameters (nutrients, climate, RFE and phenols content) as shown in Figure 2d.

To go further we moved at the community composition level and merge the three ASVs matrices (23S rRNA, *cbbL* and *bchY* genes) into a MFA (Fig.3). This MFA emphasized a clear pattern of community split between the four location (first axis) and with a second subdivision according to depth (second axis). In this MFA ordination space, community composition of Counozouls was closer to Abisko one and community composition of Männikjärve was closer to Siikaneva one (Fig.3a). Among these communities of CFMs, we found an important diversity of the microorganisms present.

**Figure 3.**
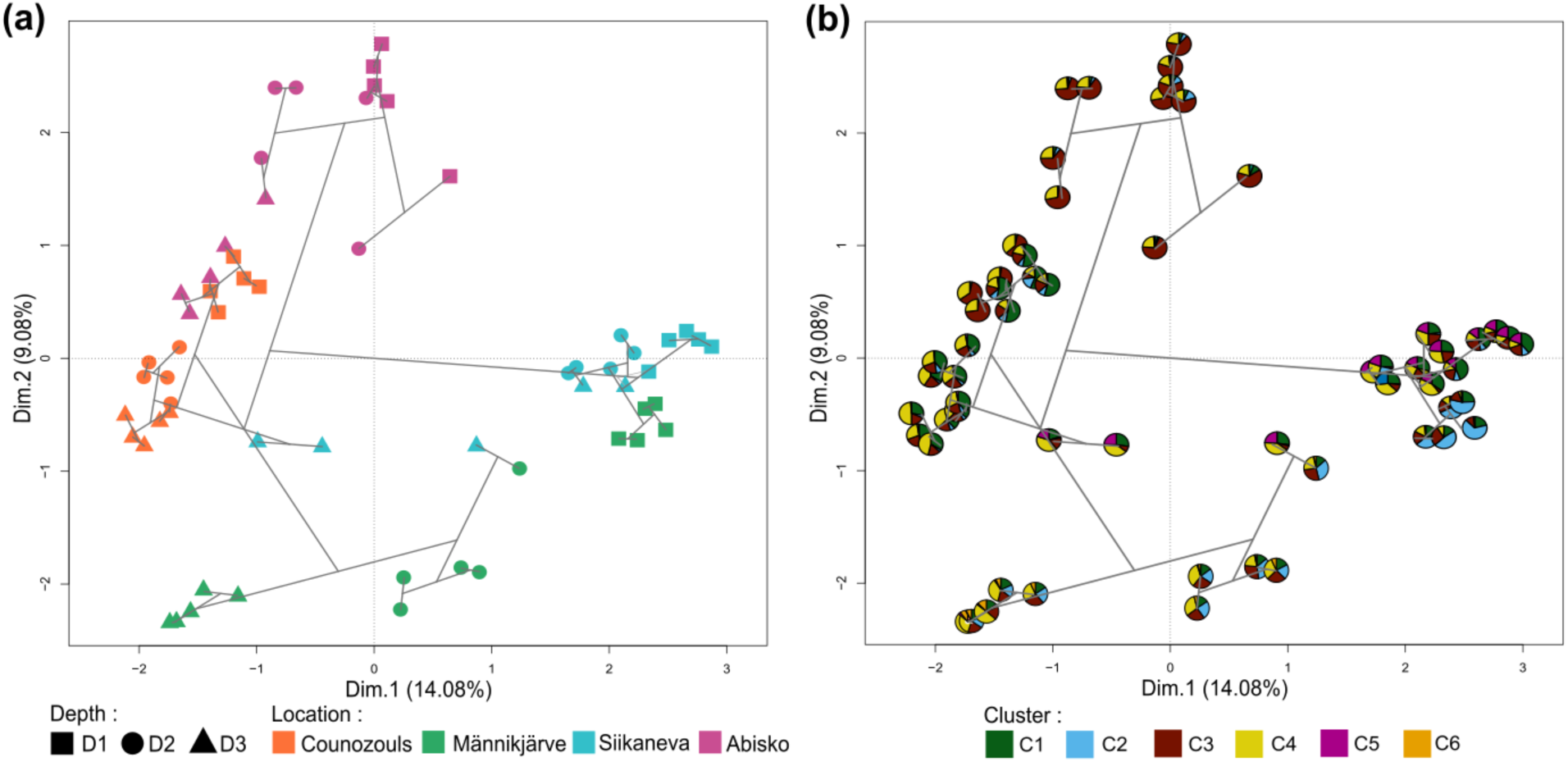
Multiple factor analysis (MFA) samples biplot of. **(a)** the 23S rRNA, *cbbL* and *bchY* genes ASVs matrices. Geometric shapes represent each sample spilt according to the four peatlands. **(b)** Pie charts represent the relative abundance of the different clusters (for the same samples) generated under the JSDM (Fig.5). Grey lines represent results of a hierarchical agglomerative clustering.

**Figure 4.**
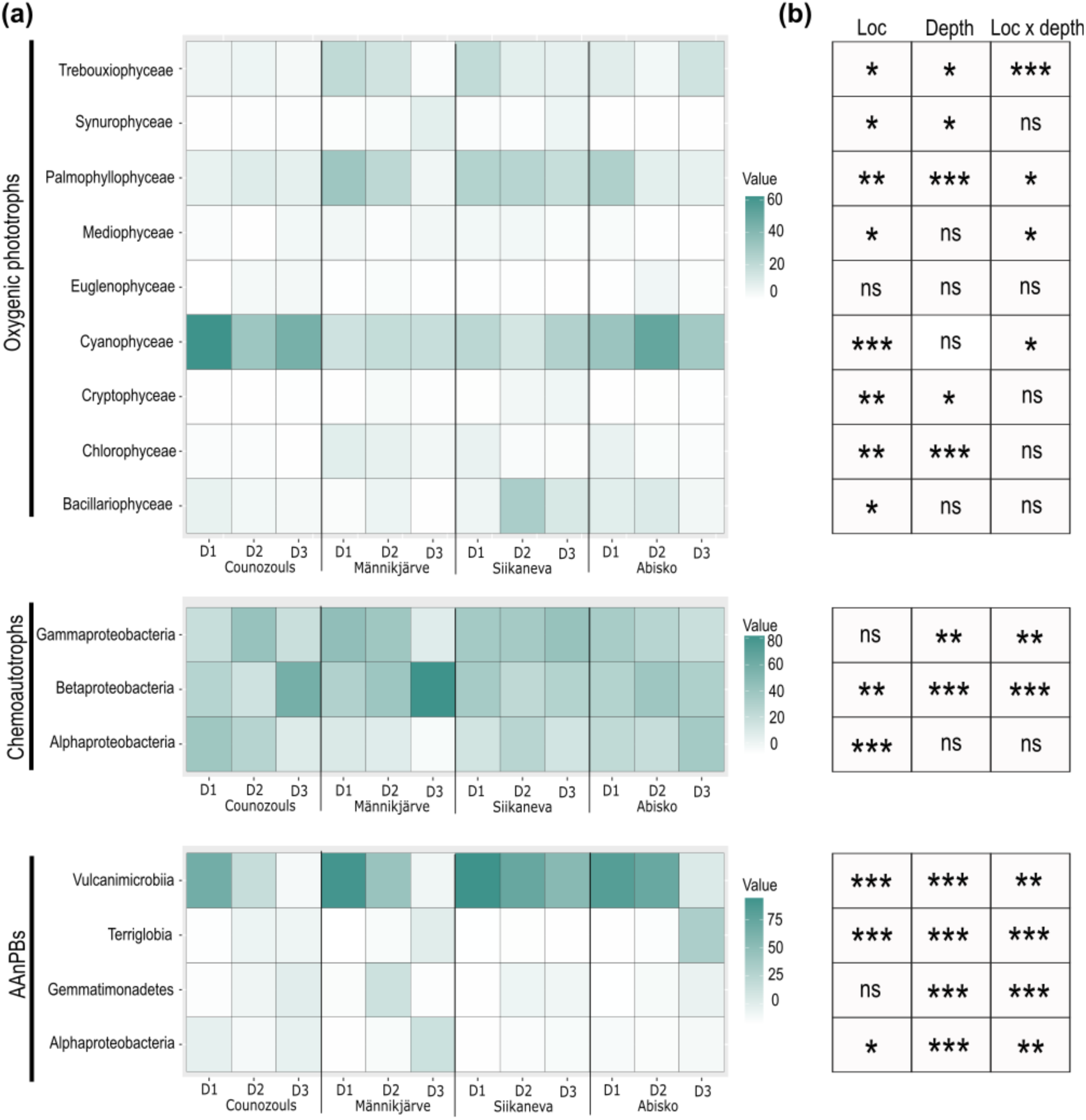
Impact of location and depth on relative abundance of ASVs aggregated by class. **(a)** Heatmaps showing the relative abundance of ASVs aggregated by class according to location and depth. Only classes with abundance higher than 5% were kept. Light red represents low abundances while dark red represents higher abundances. **(b)** P-values of the explanatory power of location, depth and location with depth. Loc = location; ns = not significant (P > 0.05); * = 0.01 < *P* < 0.05; ** = 0.001 < *P* < 0.01 and *** = *P* < 0.01; D1 = 0-5 cm; D2 = 5-10 cm and D3 = 10-15 cm.

Notably, oxygenic phototrophic ASVs belonged to 25 phyla, 41 classes, 76 orders and 102 families. These ASVs were dominated by Cyanophyceae followed by Palmophyllophyceae (Fig.4a; Fig.S11a; Fig.S12). Cyanophyceae were abundant in the four location and at all depths (Fig.4a). They were dominated by Nostocaceae and Chroococcidiopsidaceae families (Fig.S11a). Palmophyllophyceae were mostly present in the surface samples of Männikjärve, Siikaneva and Abisko (Fig.4a) and were dominated by Prasinococcaceae (Fig.S11a). We also found that Trebouxiophyceae, Bacillariophycaea, Synurophycaea and Chlorophyceae were abundant in the samples (Fig.4a). Among oxygenic phototrophs we further found three species dominating the different sites, namely *Chroococcidiopsis sp.*, *Prasinoderma coloniale* and *Anabaena sp.*. Location, depth and location together with depth had a significant impact on oxygenic phototrophs ASVs both at the class and at the family level (Fig.4b; Fig.S11b). For instance, Cyanophyceae were impacted by location (*P* < 0.001) while Palmophyllophyceae were impacted both by location and depth (*P* < 0.01; Fig.4b). However, these ASVs only represented 30% of the total 23S rRNA sequences as 60% was not assigned because the sequences were not retrieved in the database.

Chemoautotrophs ASVs belonged to 1 phylum for 3 classes, 7 orders and 13 families. Among chemoautotrophs, Proteobacteria, particulalry Beta- and Gammaproteobacteria dominated all samples (Fig.4a; Fig.S11a; Fig.S13). Betaproteobacteria were notably abundant in Counozouls and Männikjärve deepest samples (Fig.4a). Betaproteobacteria were represented by three main families, namely Thiobacillaceae, Gallionellaceae and Comamonadaceae (Fig.S11a). Gammaproteobacteria were present at all site in D1 and D2 samples (Fig.4a) with Ectothiorhodospiraceae and Acidiferrobacteraceae being the main families present (Fig.S11a). Betaproteobacteria were affected both by location and depth (*P* < 0.01) while Gammaproteobacteria were only affected by depth (*P* < 0.01; Fig.4b). Among chemoautotrophs we found *Thiobaccilus sp.*, *Nitrobacter winogradsky*, *Sulfurifustis variabilis*, *Sulfuricaulis limicola* and *Hydrogenophaga sp*. being the most abundant species.

AAnPBs ASVs were composed of 11 phyla, 20 classes, 28 orders and 35 families among which Terriglobia, Gemmatimonadetes, Alphaproteobacteria and Vulcanimicrobiia were the most abundant classes with a net dominance of Vulcanimicrobiia for depth D1 and D2 (Fig.4a; FigS11a; Fig.S13). At the family level, the family Vulcanimicrobiaceae also dominated largely (Fig.S11a). Vulcanimicrobiia were significantly affected by both location (*P* < 0.001) and depth (*P* < 0.001; Fig.4b).

#### CFMs communities group in distinct clusters

We identified six major clusters of CFMs with shared within-cluster responses, but opposite between-cluster responses, to environmental conditions (Fig. 5a). Each cluster was dominated by chemoautotrophs, except clusters 4 and 6 which were dominated by oxygenic phototrophs (Fig. 5b). Each cluster was also characterized by CFMs with different niche size (Fig.5c). Clusters 3 and 4 exhibited ASVs with the largest niche size (Fig.5c) and were logically widespread across all sites and depths (Fig.3b). These clusters were both characterized by high abundances of oxygenic phototrophs (Fig.5b), including Palmophyllocyceae, Cyanophyceae and Baccillariophyceae (Fig.S15a and Fig.S16a), and chemoautotrophs (Fig.5b) with Proteobacteria, mainly Acidiferrobacteraceae, Nitobacteraceae and Ectothiorhodospiraceae (Fig.S15b and Fig.S16b). These clusters responded differently to environmental conditions with cluster 3 related to cold and wet conditions and high concentrations of phosphorus and nitrogen whilst cluster 4 was driven by dry conditions and an environment rich in carbon and phosphorous (Fig.5e).

**Figure 5.**
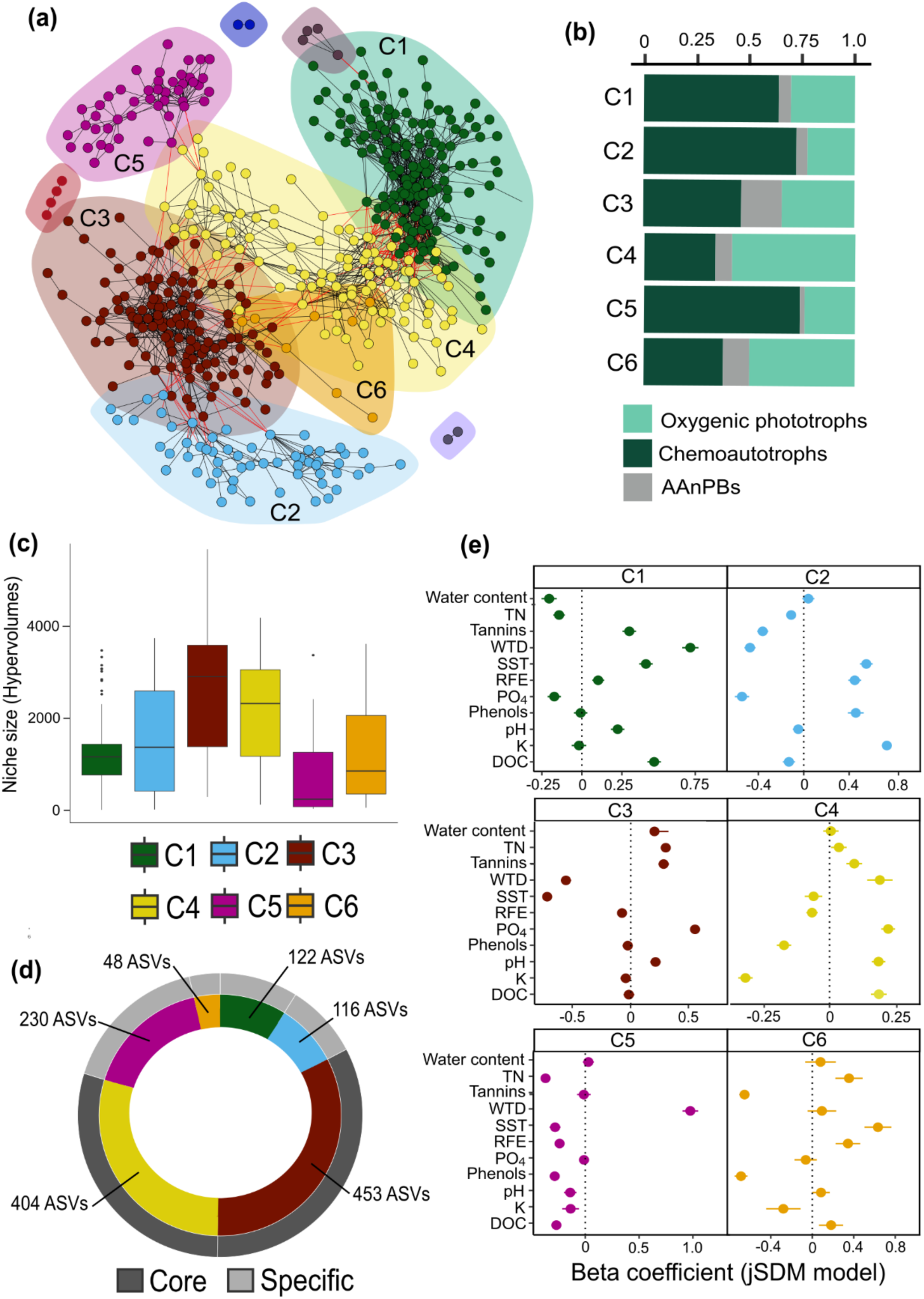
**Co-occurrence network of the variation between ASVs caused by environmental parameters**. **(a)** Clusters of co-occurring ASVs with each dot representing one ASVs. **(b)** Barplot of the relative abundance of each microbial group (*cbbL* gene – chemoautotrophs; *bchY* gene – AAnPBs and 23S rRNA gene – oxygenic phototrophs) within each cluster. **(c)** Niche size (hypervolumes) of each cluster. **(d)** Absolute distribution of ASVs into the six clusters and according if these ASVs are considered core microbiome ASVs (dark grey) or specific microbiome ASVs (light grey). **(e)** Mean of beta coefficient from the jSDM model showing how environmental parameters are affecting the clusters. C1 = cluster 1; C2 = cluster 2; C3 = cluster 3; C4 = cluster 4; C5 = cluster 5; C6 = cluster 6; TN = total nitrogen; DOC = dissolved organic carbon; SST = spring soil temperature; WTD = water table depth; D1 = 0-5 cm; D2 = 5-10 cm and D3 = 10-15 cm.

Clusters 1, 2 and 6 showed intermediate niche sizes (Fig.5c). They were accordingly not present in all sites and depths (Fig.3b). Clusters 1 and 2 were dominated by chemoautotrophs (Fig.5b) with a lot of Alpha and Betaproteobacteria among which Nitrobacteraceae, Thiobacillaceae and Comamonadaceae dominated (Fig.S15b and Fig.S16b) followed by oxygenic phototrophs (Fig.5b) composed of diverse classes and families (Fig.S15a and Fig.S16a). Cluster 6 was composed of unclassified chemoautotrophs and Beta and Gammaproteobacteria (Fig.S15a, b, c and Fig.S16 a, b and c). Clusters 1 and 6 were associated with warm, C and N-rich conditions. Cluster 2 was associated with wet and warm conditions, as well as with dissolved organic matter rich in phenols and samples rich in potassium (Fig.5e). Cluster 5 had the lowest hypervolumes (Fig.5c) and was present only in a few samples (Fig.3b), mainly in Siikaneva. This cluster was dominated by chemoautotrophs (Fig.5b) with Betaproteobacteria (Thiobacillaceae and Comemonadaceae; Fig.S15b and Fig.S16b) and by some specific oxygenic phototrophs (Fig.S15a and Fig.S16a) and was strongly related to dry and nutrient-poor conditions (Fig.5e).

Following the results of niche size we further classified ASVs belonging to clusters 3 and 4 as the core microbiome present in all our samples (Fig.5d) and ASVs belonging to clusters 1, 2, 5 and 6 as the specific ASVs only present in a few sites. Clusters 3 and 4 shared 62% of the ASVs (453 ASVs – 33% for C3 and 404 ASVs – 29% for C4) whilst Cluster 1 represented 9% (122 ASVs), Cluster 2, 8% (116 ASVs), Cluster 5, 17% (230 ASVs) and Cluster 6 only 4% (48 ASVs) of the ASVs.

### Drivers of CFM abundance

We used structuration equation modelling (SEM) to decipher the mechanisms underlying CFM abundance in peatlands (Fig.6). Our SEM revealed that total CFM abundance — a latent variable defined by the absolute quantification of the 23S rRNA gene (path = 0.82), of the *cbbL* gene (path = 0.87), and of the *pufM* gene (path = 1.02) — was directly mediated by changes in environmental conditions (path = 1.03). The environment also indirectly influenced CFM total abundance through changes in CFM community structures. Indeed, changing environmental conditions directly influenced the relative abundance of each cluster (cluster 1 path = 0.66, cluster 2 path = 0.37 and cluster 3 path = -0.64), as well as CFM richness (Chao1, path = 0.61) and community structure (MFA1, path = -1.43). Then, the relative abundance of cluster 1 (path = 0.68) and 3 (path = -0.43) influenced the community structure of all CFMs (MFA1), while the relative abundance of cluster 2 influenced both Chao1 and MFA1 (path = -0.57 and path = 0.84, respectively) and the relative abundance of cluster 3 influenced MFA1 (path = -0.43), which in turn determined CFM abundance (Chao1, path = 0.89; Cluster 3, path = 0.67).

**Figure 6.**
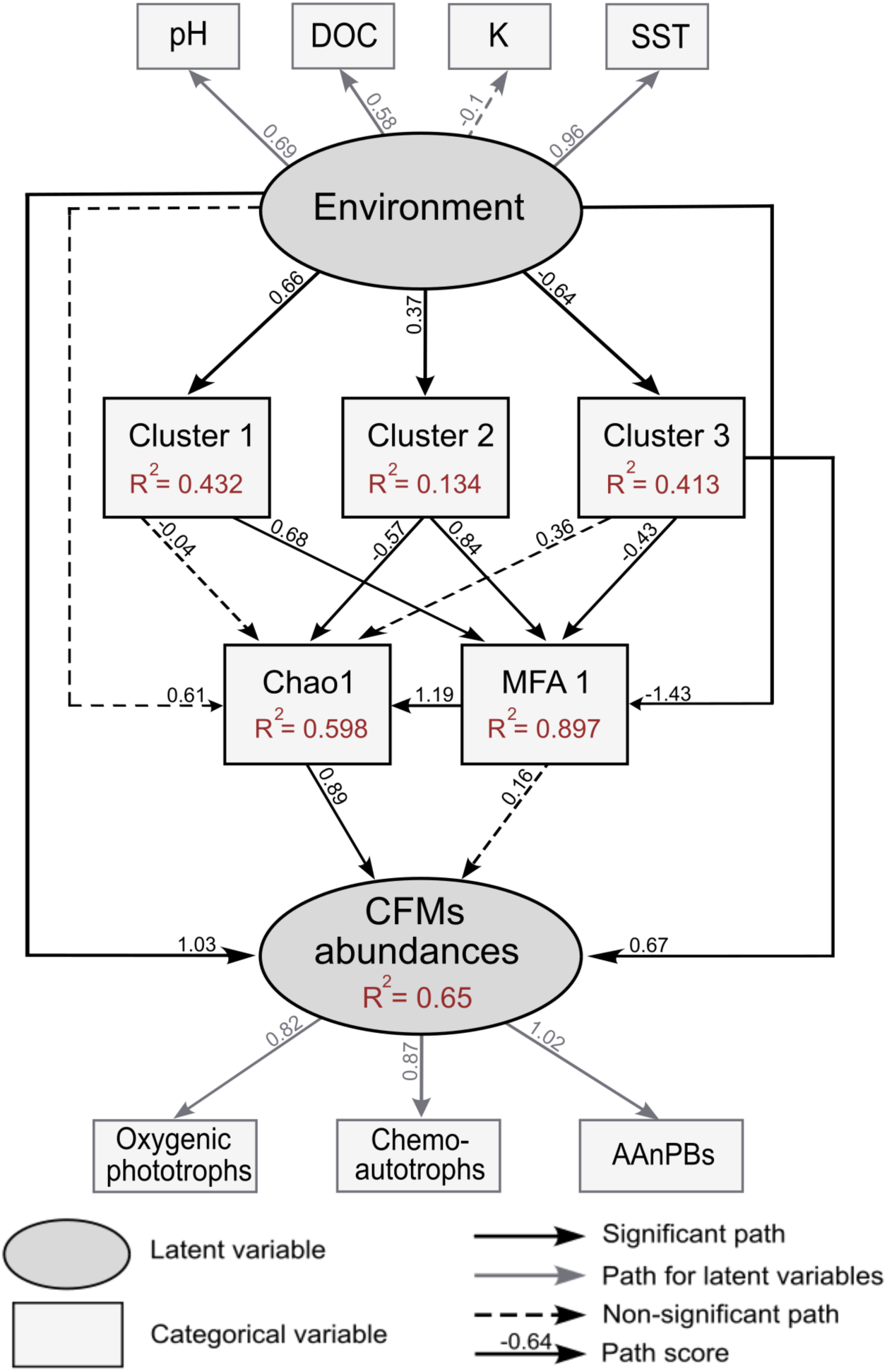
Drivers and mechanisms behind absolute quantification of carbon fixing microorganisms (CFMs) in peatlands. Structural equation model (SEM) of the correlation between environmental parameters, clusters relative abundances, richness (Chao1), community structure (MFA 1) and CFMs gene absolute quantification. Adjusted R-squared in the boxes indicate the percentage of variance explained by each model. Numbers along the arrows indicate the weight of the path relationship. Dotted arrows indicate non-significant (*P* > 0.05) relationship. SST = spring soil temperature; DOC = dissolved organic carbon. *Environment* and *CFMs abundances* are two latent variables defined by pH, DOC, K and SST for *Environment* and by the abundance of 23S (oxygenic phototrophs), *cbbL* (chemoautotrophs) and *pufM* (AAnPBs) genes for *CFMs abundances*.

## Discussion

Our aim was to explore CFMs abundance, diversity and community composition as well as to identify their main environmental drivers. We found that CFMs were abundant regardless the depth or the peatland type. With 1.15 x 10^8^ ± 1.2 x 10^7^ copies g^-1^ dw, CFMs represented ∼40% of the total bacterial abundance (Fig.2a). This corroborates recent observations (Le Geay et al., 2024b), and highlight the potential of CFMs for carbon fixation in peatlands through the CBB cycle. This high abundance was supported by a strong diversity spanning 37 phyla for 7,960 ASVs. However, this diversity was not consistent among all sites. We found that CFMs were structured into different communities that were either present in a few site or widespread across all sites in relation to environmental conditions. This highlighted the presence of a core microbiome that was modulating CFM diversity and abundances across sites. We caution that we only focused on CFMs utilizing the CBB cycle while many other pathways exist (Berg, 2011). Nevertheless, this is the first demonstration of the structuration and abundance of different CFMs communities in peatlands, which lays the foundation to better understand the role of microorganisms in peatland primary productivity and how they could contribute to microbial carbon sinks.

### CFMs are abundant in northern peatlands

Our findings reveal that although peatlands exhibit very specific environmental constraints, they show very similar CFM abundances. Indeed, both oxygenic phototrophs and chemoautotrophs abundances were comparable to those found in deserts, forests, grasslands, or wetlands soils with around 10^5^ - 10^8^ copies.g^-1^ of dry soil (Li et al., 2013; Mar Lynn et al., 2017; Sauze et al., 2017; Jassey et al., 2022; Liao et al., 2023). AAnPBs abundance was also similar to those found in soils (10^5^ - 10^9^ copies.g^-1^ of dry soil ; Feng et al., 2011b, 2011a; Sato-Takabe et al., 2016, 2020). However, AAnPBs were more abundant in peatlands than in aquatic systems which previously showed low AAnPBs abundances ranging from 10^2^ to 10^5^ cells.mL^-1^ (Lamy et al., 2011; Ritchie and Johnson, 2012). We further expected depth to have an important impact on these abundances as oxygenic phototrophs and AAnPBs are sensitive to the presence of light and oxygen, that are mostly available in the first top centimeters (Bengtsson et al., 2018; Hädrich et al., 2019), while chemoautotrophs are linked to the presence of reduced compounds more concentrated in deeper layers (Rydin and Jeglum, 2006; Hädrich et al., 2019). If we saw a decrease of phototrophs with depth, we did not see a real increase of chemoautotrophs with depth meaning we may not have sample deed enough.

This important abundance could support CFM driven C fixation as microbial abundance is usually related to metabolic activity (Morales et al., 2010; Delgado-Baquerizo et al., 2016a; Hamard et al., 2021a; Sáez-Sandino et al., 2023). In particular, chemoautotrophs have been shown to perform C fixation in peatlands (Gilbert et al., 1998) and Bay et a., 2021 showed that genes abundances were related to C fixation rates. Taken together, these results suggest that chemoautotrophs need to be considered for the contribution to peatlands primary productivity and that previous estimation only based on oxygenic phototrophs (Hamard et al., 2021a) probably greatly underestimate C fixation conducted by microorganisms. Moreover, it is important to note that we only focused on the CBB cycle while CFMs are diverse and using other pathways such as the Wood–Ljungdahl pathway that can be energetically favorable (Berg, 2011) which increases even more the C fixation potential performed by CFMs.

#### Biodiversity patterns

The high CFM abundance was supported by a high biodiversity, with on average 312 oxygenic phototrophs, 563 chemoautotrophs, and 146 AAnPBs. We found CFM classes that are commonly found in other soils, such as Cyanophyceae, Palmophyllophyceae, Bacillariophyceae, Synurophyceae or Trebouxiophyceae for oxygenic phototrophs and Nitrobacteraceae, Thiobacillaceae and Acidiferrobacteraceae for chemoautotrophs (Cano-Díaz et al., 2020; Oliverio et al., 2020; Feng et al., 2022; Jassey et al., 2022; Li et al., 2023). For AAnPBs we found one phylum, Vulcanimicrobiota also known as “Candidatus Eremiobacterota”, completely dominating the samples that is not usually dominating. This phylum has also been found in permafrost soils (Woodcroft et al., 2018), boreal mosses (Holland-Moritz et al., 2018; Ward et al., 2019) and other peatlands (Serkebaeva et al., 2013). Bacteria of the phylum Vulcanimicrobiota are metabolically diverse and versatile, performing photoautotrophy (Holland-Moritz et al., 2018; Ward et al., 2019) and chemolitoautotrophy (Ray et al., 2020; Ji et al., 2021) allowing them to adapt to acidic, contaminated as well as nutrient poor environment (Ward et al., 2019; Ji et al., 2021). Recently, Yabe et al., (2022) succeeded for the first time in the isolation of a “Candidatus Eremiobacterota” representative, *Vulcanimicrobium alpinus.* This species demonstrated bacteriochlorophyll biosynthesis, CO_2_ fixation and phototrophic motility capabilities. Thus, the presence of Vulcanimicrobia in peat samples suggests that AAnPBs can fix CO_2_ in peatlands. As showed by Yabe et al., (2022), these AAnpBs may not rely on CO_2_ fixation for the CBB cycle ; instead, they might utilize anaplerotic pathway to replenish the citric-acid cycle to maintain optimal functioning under nutrient poor conditions.

#### A CFM core microbiome

We found that CFM diversity can be structured into six main clusters, all shaped by specific, and sometimes antagonistic, responses to environmental conditions (Fig. 5). These clusters showed different ecological niche size (Fig.5 ; Blonder, 2018), meaning that some clusters were ubiquitous (C3 and C4) while others were only present in a few site (C1, C2, C5 and C6). Taken together, these findings evidenced a core (C3 and C4) and a more specific CFM microbiome (C1, C2, C5 and C6). The core microbiome was strongly related to temperature, WTD, pH, C and P. It was composed of a mix of families belonging to the three CFMs groups. The most abundant species retrieved were *Nitrobacter winogradsky* and *N. vulgaris*, which are important players of the C and N cycles (Starkenburg et al., 2006), *Anabaena sp.* a common carbon and nitrogen fixing cyanobacteria (Hrouzek et al., 2004), *Prasinoderma coloniale*, a green alga adapted to low light and oligotrophic habitat (Li et al., 2020) and Vulcanimicrobiia (AAnPBs). Species able to utilizing H_2_ oxidation to derive energy (Janssen et al., 2010) like *Hydrogenophaga sp*. and *Cupriavidus metallireducens*, and species able to oxidize thiosulfate, tetrathionate as well as elemental sulfur (Kojima et al., 2015, 2016) such as *Sulfurisistis variabilis* and *Sulfuricaulis limicolica* were also found in this core microbiome. The specific microbiome greatly differs from the core microbiome, with an important species turnover between clusters and specific responses to local environmental conditions such as temperature, WTD and nutrient availability. ASVs composing the specific CFM microbiome essentially belonged to Burkholderiaceae, Nitrobacteraceae and Thiobacillaceae families with two species more abundant, *Thiomonas sp.*, adapted to low pH and low concentration of nutrients (Arsène-Ploetze et al., 2010) and *Thiobaccillus sp.*, that fix CO_2_ through the oxidation of inorganic sulfur compounds (Skinner and Jahren, 2007). Some very specific species were also retrieved such as *Acidihalobacter prosperus*, a chemoautotroph bacteria acidophile and halotolerant able to use either ferrous iron or reduced sulfur as electron donor (Khaleque et al., 2020) to fix CO_2_ and *Chaetosphaeridium globosum*, a freshwater green algeae (Gibson, 2008) that were only present in cluster 5.

We found that the core microbiome was an important driver of CFMs richness, diversity, community structure. Its response to environmental conditions also strongly modulated CFMs total abundances. Notably, temperature, pH and C content, known to drive communities structure (Delgado-Baquerizo et al., 2016b; Oliverio et al., 2020; Jassey et al., 2022), shaped CFMs communities. For instance, clusters 3 responded negatively to the environment while clusters 1 and 2 responded positively (Fig.5 and 6) meaning that even small changes in the environment can re-structure these communities and have important feedback for the peatland C cycle.

#### General implications

Carbon fixation in ecosystems is underpinned by at least seven biochemical pathways, of which six are exclusively found among bacteria and archaea (Berg, 2011). This study provide a comprehensive picture on how three major microbial carbon fixation pathways are distributed among peatlands and discover the unexpected abundance and occurrence of certain pathways and taxonomic groups. In particular, we evidenced the importance of the core CFM microbiome in modulating CFM diversity and abundance across environmental gradients. We caution that our study is DNA-based and could therefore not demonstrate potential activities or functions as DNA is only given a description of the community at a given moment. As such, further studies are needed to link our results to functionality, by using RNA in completement to DNA or by measuring CFM CO_2_ flux rates (Urich et al., 2008). Nevertheless, these findings have significant implications as they show that much greater diversity of bacteria and archaea have the potential to contribute to peatland carbon fiation than previously expected. As CFMs as the potential to mitigate to some extent microbial C emission under climate change in peatlands (Le Geay et al., 2024a) our findings illustrate the importance of considering not only oxygenic phototrophs but also chemoautotrophs and AAnPBs in peatland C fixation.

## Acknowledgments

This work has been supported by the MIXOPEAT (Grant No. ANR-17-CE01-0007 to VEJJ) and BALANCE (Grant No. ANR-23-ERCC-0001-01 to VEJJ) projects funded by the French National Research Agency. This study has been partially supported through the grant EUR TESS (Grant No. ANR-18-EURE-0018 to MLG) in the framework of the Programme des Investissements d’Avenir. We thank the Plateforme Analyses Physico-Chimiques from the Centre de Recherche sur la Biodiversité et l’Environnement (Université de Toulouse, France). We thank the Swedish Polar Research Secretariat and SITES for the support of the work done at the Abisko Scientific Research Station. SITES is supported by the Swedish Research Council’s grant 4.3-2021-00164. We are grateful to Lisa Vayssier for her help in the laboratory.

## Data Accessibility and Benefit-Sharing

The datasets generated during and/or analysed during the current study are available in the [NAME] repository, [PERSISTENT WEB LINK TO DATASETS].

## Author contribution

MLG, BL and VEJJ conceived the project. MLG, VEJJ, and MK collected the samples. MLG performed laboratory work and analyses under the supervision of VEJJ, BL and KM. MLG performed data analyses with the help of VEJJ. MLG wrote the original draft and MLG and VEJJ led the writing of the manuscript with input from all other authors. All author reviewed and edited the manuscript.

## Conflict of interest statement

The authors declare no conflicts of interest

## Notes

### Competing Interest Statement

The authors have declared no competing interest.

