## Supplementary figures and tables for "Diversity, abundance, and biogeography of CO_2_ fixing microorganisms in peatlands"

### Diversity and abundance of $CO_2$ fixing micro-organisms in peatlands

#### Table of Contents:


| | |
|---|---|
| Table S1. Primer pairs used in this study<br>Table S2. PCR reaction conditions used in this study. | Page 3 |
| Table S3. Reaction conditions for ddPCR | Page 4 |
| Table S4. Environmental and chemicals characterization of each site<br>Table S5. Results of ANOVA testing the impact of location and depth on different environmental parameters including metabolites, organic matter quality, nutrients and climatic conditions. | Page 5 |
| Table S6. Results of posthoc test (after ANOVA) testing the impact of location and depth on different environmental parameters including metabolites, organic matter quality, nutrients and climatic conditions. | Page 6 |
| Table S7. Results of ANOVA testing the impact of location and depth on gene abundance.<br>Table S8. Results of posthoc test (after ANOVA) testing the impact of location and depth on gene abundance.<br>Table S9. Results of ANOVA testing the impact of location and depth on 23S rRNA gene and *bchY* gene alpha richness (Chao1 index) and alpha diversity (Shannon index). | Page 8 |
| Table S10. Results of posthoc test (after ANOVA) testing the impact of location and depth on 23S rRNA gene, *cbbL* gene and *bchY* gene richness (chao1 index) and alpha diversity (Shannon index). | Page 9 |
| Figure S1. Rarefaction curves | Page 10 |
| Figure S2. Correlation plot of environmental variables | Page 11 |
| Figure S3. Boxplot describing (a) pH, (b) DOC, (c) TN and (d) WTD for each location<br>Figure S4. Boxplot describing organic matter quality index (DOCq, peak_A, peak_C, peak_M, RFE, freshness, BIX and FI) for each location | Page 12 |
| Figure S5. Boxplot describing cations ($Na^+$, $NH_4^+$, $K^+$, $Mg^{2+}$ and $Ca^{2+}$) and anions ($F^-$ $NO_3^-$, $Br^-$, $SO_4^{2-}$ and $PO_4^{3-}$) for each location | Page 13 |
| Figure S6. Boxplot describing metabolites (carbohydrates, flavonoids, phenols, tannins and water phenols) for each location | Page 14 |
| Figure S7. Absolute quantification using ddPCR | Page 15 |

| | |
|---|---|
| Figure S8. Absolute quantification of cyanobacteria using ddPCR | Page 16 |
| Figure S9. Diversity description | Page 17 |
| Figure S10. NMDS showing the community structure | Page 18 |
| Figure S11. Impact of location and depth on relative abundance of ASVs aggregated by family | Page 19 |
| Figure S12. Upset plot of the presence of ASVs aggregated by class for the 23S rRNA gene in the four peatland sites<br>Figure S13. Upset plot of the presence of ASVs aggregated by class for the *cbbL* gene in the four peatland sites | Page 20 |
| Figure S14. Upset plot of the presence of ASVs aggregated by class for the *bchY* gene in the four peatland sites | Page 21 |
| Figure S15. Barplot of the relative abundance of each class constituting the different clusters | Page 22 |
| Figure S16. Barplot of the relative abundance of each family constituting the different clusters | Page 23 |

**Table S1.** Primer pairs used in this study

| Analysis | Microorganism targeted | Primers | Amplicon Size | Primer sequence | Reference |
|---|---|---|---|---|---|
| ***(a) PCR*** | Prokaryote | PCR1-515F | 412 bp | 5' GTG YCA GCM GCC GCG GTA 3' | (Wang & Qian, 2009) |
| | | PCR1-909R | | 5' CCC CGY CAA TTC MTT TRA GT 3' | |
| | Photosynthetic microorganisms | P23SrV_f1 | 410 bp | 5' GGA CAG AAA GAC CCT ATG AA 3' | (Sherwood & Presting, 2007) |
| | | P23SrV_r1 | | 5' CAG CCT GTT ATC CCT AGA G 3' | |
| | Chemoautotrophs | cbbL-IA-CHEM | 479 bp | 5' GAR GGN TCN GTN GTY AAC GT 3' | (Alfreider & Bogensperger, 2018b) |
| | | cbbL-IA-r | | 5' GTA RTC GTG CAT GAT GAT SGG 3' | |
| | AAnPB | bchY-fwd | 500 bp | 5' CCN CAR CAN ATG TGY CCN GCN TTY GG 3' | (Yutin et al., 2009) |
| | | bchY-rev | | 5' GGR TCN RCN GGR AAN ATY TCN CC 3' | |
| ***(b) ddPCR*** | Prokaryote | L/Prba338f | 180 bp | 5′- ACT CCT ACG GGA GGC AGC AG -3′ | (Øvreås et al., 1997) |
| | | K/Prun518r | | 5′- ATT ACC GCG GCT GCT GG -3′ | |
| | Cyanobacteria | 16SCF | 164 bp | 5'-GGC AGC AGT GGG GAA TTT TC-3' | (Oh et al., 2012) |
| | | 16SUR | | 5'-GTM TTA CCG CGG CTG CTG G-3' | |
| | Photosynthetic microorganisms | 23S255f | 160 bp | 5'- GGA TTA GAT ACC CYD GTA GTC C -3' | (Le Geay et al., 2024) |
| | | P23SrV_r1 | | 5' – TCA GCC TGT TAT CCC TAG AG -3' | (Sherwood & Presting, 2007) |
| | Chemoautotrophs | cbbLR1F | 274 bp | 5'- AAG GAY GAC GAG AAC ATC -3' | (Selesi et al., 2007) |
| | | cbbLRintR | | 5'- TGC AGS ATC ATG TCR TT -3' | |
| | AAnPB | pufM forward 557 | 193 bp | 5'-TAC GGS AAC CTG TWC TAC-3' | (Du et al., 2006) |
| | | pufM reverse 750 | | 5'-CCA TSG TCC AGC GCC AGA A-3' | |

**Table S2.** CR reaction conditions used in this study.

| Targeted gene | 16S rRNA | 23S rRNA | *bchY* | *cbbL* |
|---|---|---|---|---|
| Annealing temperature | 55˚C | 55˚C | 50˚C | 52˚C |
| PCR conditions | 95°C – 10 min<br><br>35 cycles :<br>94°C – 60 sec<br>55°C – 40 sec<br>72 ˚C – 30 sec<br><br>72°C – 10 min | 95°C – 10 min<br><br>35 cycles :<br>94°C – 60 sec<br>55°C – 45 sec<br>72 ˚C – 45 sec<br><br>72°C – 10 min | 95°C – 10 min<br><br>45 cycles :<br>94°C – 60 sec<br>50°C – 45 sec<br>72 ˚C – 60 sec<br><br>72°C – 10 min | 95°C – 10 min<br><br>38 cycles :<br>94°C – 60 sec<br>52°C – 30 sec<br>72 ˚C – 60 sec<br><br>72°C – 10 min |

**Table S3.** Reaction conditions for ddPCR

| Primers | 16SCF/<br>16SUR | L/Prba338f/<br>K/Prun518r | 23S_new/<br>P23Srv-r1 | pufMfwd557/<br>pufMrev750 | cbbLR1F/<br>cbbLRintR1 |
|---|---|---|---|---|---|
| Primers concentration | 150 nM | 250 nM | 250 nM | 250 nM | 250 nM |
| DNA dilution | 1/100 | 1/1,000 | 1/100 (D1, D2) and 1/10 (D3) | 1/100 (D1, D2) and 1/10 (D3) | 1/10 |
| Annealing temperature | 52˚C | 61˚C | 57.6˚C | 50.2˚C | 53˚C |
| PCR conditions | 95°C – 5 min<br><br>40 cycles :<br>94°C – 30 sec<br>52°C – 60 sec<br><br>4°C – 5 min<br>98°C – 10 min<br>Ramp rate = 2°C/sec | 95°C – 5 min<br><br>40 cycles :<br>94°C – 30 sec<br>61°C – 30 sec<br><br>4°C – 5 min<br>98°C – 10 min<br>Ramp rate = 1°C/sec | 95°C – 5 min<br><br>40 cycles :<br>94°C – 30 sec<br>57.6°C – 60 sec<br><br>4°C – 5 min<br>98°C – 10 min<br>Ramp rate = 2°C/sec | 95°C – 5min<br><br>45 cycles :<br>94°C – 60 sec<br>50.2°C – 2 min<br><br>4°C – 5 min<br>98°C – 10 min<br>Ramp rate = 1°C/sec | 95°C – 5min<br><br>45 cycles :<br>94°C – 60 sec<br>53°C – 2 min<br><br>4°C – 5 min<br>98°C – 10 min<br>Ramp rate = 1°C/sec |

**Table S4. Environmental and chemicals characterization of each site.** Cou = Counozouls; Män = Männikjärve; Siik = Siikaneva; Abi = Abisko; DOC = dissolved organic carbon; TN = total nitrogen; WAT = winter air temperature; WST = winter soil temperature; WP = winter precipitation; WWTD = winter water table depth; SAT = spring air temperature; SST = spring soil temperature; SP = spring precipitation and SWTD = spring water table depth.

| | pH | DOC | TN | WAT | WST | WP | WWTD | SAT | SST | SP | SWTD |
|---|---|---|---|---|---|---|---|---|---|---|---|
| **Cou** | 5.8 | 83.8 | 1.8 | -1.3 | 1.4 | 85.0 | 462.0 | 6.4 | 7.1 | 275.0 | 472.1 |
| **Män** | 5.0 | 44.4 | 1.3 | -4.7 | 0.9 | 105.0 | 358.6 | 5.5 | 5.5 | 142.0 | 391.3 |
| **Siik** | 4.9 | 32.5 | 0.7 | -6.0 | 0.2 | 0.0 | 681.2 | 2.5 | 3.1 | 177.0 | 768.2 |
| **Abi** | 4.7 | 52.3 | 1.4 | -10.2 | -0.6 | 5.0 | 40.8 | 2.9 | 2.0 | 26.0 | 114.5 |

**Table S5. Results of ANOVA testing the impact of location and depth on different environmental parameters including metabolites, organic matter quality, nutrients and climatic conditions.** DOC = dissolved organic carbon, TN = total nitrogen, WAT = winter air temperature, WST = winter soil temperature, WP = winter precipitation, WWTD = winter water table depth, WPAR = winter PAR, SAT = spring air temperature, SST = spring soil temperature, SP = spring precipitation, SWTD = spring water table depth, SPAR = spring PAR, Cou = Counozouls, Män = Männikjärve, Siik = Siikaneva. Abi = Abisko, D1 = 0-5 cm, D2 = 5-10 cm, D3 = 10-15 cm. Bold results mean significant *p-value*.

| | Location | | Depth Cou | | Depth Män | | Depth Siik | | Depth Abi | |
|---|---|---|---|---|---|---|---|---|---|---|
| | *F-value* | *p-value* | *F-value* | *p-value* | *F-value* | *p-value* | *F-value* | *p-value* | *F-value* | *p-value* |
| DOCq | 3.325 | **0.026** | - | - | - | - | - | - | - | - |
| Peak_A | 17.27 | **<0.001** | - | - | - | - | - | - | - | - |
| Peak_C | 15.9 | **<0.001** | - | - | - | - | - | - | - | - |
| Peak_M | 14.42 | **<0.001** | - | - | - | - | - | - | - | - |
| RFE | 18.73 | **<0.001** | - | - | - | - | - | - | - | - |
| Freshness | 64.57 | **<0.001** | - | - | - | - | - | - | - | - |
| BIX | 40.02 | **<0.001** | - | - | - | - | - | - | - | - |
| FI | 10.81 | **<0.001** | - | - | - | - | - | - | - | - |
| WTD | 253.9 | **<0.001** | - | - | - | - | - | - | - | - |
| Tannins | 4.387 | **0.008** | 1.674 | 0.228 | 14 | **<0.001** | 2.591 | 0.116 | 1.794 | 0.208 |
| Water_Phenols | 0.446 | 0.721 | 0.148 | 0.864 | 2.42 | 0.131 | 3.117 | 0.081 | 0.475 | 0.633 |
| Carbohydrates | 6.11 | **0.001** | 1.787 | 0.209 | 8.036 | **0.006** | 0.447 | 0.65 | 2.107 | 0.164 |
| Flavonoids | 0.216 | 0.885 | 0.504 | 0.616 | 4.469 | **0.035** | 0.776 | 0.482 | 0.88 | 0.44 |
| Phenols | 1.5 | 0.225 | 1.741 | 0.217 | 7.925 | **0.006** | 1.085 | 0.369 | 0.282 | 0.759 |
| $Na^+$ | 0.35 | 0.79 | 3.46 | 0.065 | 0.418 | 0.668 | 1.12 | 0.358 | 0.227 | 0.8 |
| $NH_4^+$ | 1.747 | 0.168 | 1 | 0.397 | 0.685 | 0.523 | 1.041 | 0.383 | 0.396 | 0.699 |
| $K^+$ | 1.673 | 0.183 | 8.123 | **<0.001** | 10.25 | **0.0025** | 24.8 | **<0.001** | 5.6255 | 0.019 |
| $Mg^{2+}$ | 15.44 | **<0.001** | 0.501 | 0.618 | 1.018 | 0.391 | **0.044** | 0.957 | 1.063 | 0.376 |
| $Ca^{2+}$ | 1.81 | 0.156 | 1.305 | 0.307 | 1.783 | 0.21 | 0.72 | 0.507 | 0.666 | 0.532 |
| $F^-$ | 2.858 | **0.045** | 6.23 | 0.0139 | 0.423 | 0.659 | 0.882 | 0.439 | 0.97 | 0.407 |
| $CL^-$ | 1.233 | 0.306 | 0.672 | 0.529 | 0.415 | 0.669 | 0.445 | 0.651 | 0.645 | 0.542 |
| $NO_2^-$ | 0.844 | 0.476 | 2.209 | 0.152 | 0.474 | 0.634 | 0.075 | 0.928 | 0.483 | 0.629 |

| | | | | | | | | | | |
|---|---|---|---|---|---|---|---|---|---|---|
| $NO_3^-$ | 2.81 | **0.0476** | 1.519 | 0.258 | 5.058 | **0.026** | 1.624 | 0.238 | 0.277 | 0.763 |
| $BR^-$ | 4.194 | **<0.001** | 4.33 | **0.038** | 2.863 | 0.096 | 3.5 | 0.0635 | 11.81 | 0.002 |
| $SO_4^{2-}$ | 1.36 | 0.264 | 0.159 | 0.855 | 0.288 | 0.755 | 1.26 | 0.319 | 0.149 | 0.863 |
| $PO_4^{3-}$ | 32.67 | **<0.001** | 1.281 | 0.313 | 1.22 | 0.329 | 0.761 | 0.489 | 1.616 | 0.239 |
| PH | 24.82 | **<0.001** | - | - | - | - | - | - | - | - |
| DOC | 36.33 | **<0.001** | - | - | - | - | - | - | - | - |
| TN | 26.64 | **<0.001** | - | - | - | - | - | - | - | - |
| WAT | >100 | **<0.001** | - | - | - | - | - | - | - | - |
| WP | >100 | **<0.001** | - | - | - | - | - | - | - | - |
| WST | >100 | **<0.001** | - | - | - | - | - | - | - | - |
| WPAR | >100 | **<0.001** | - | - | - | - | - | - | - | - |
| WWTD | >100 | **<0.001** | - | - | - | - | - | - | - | - |
| SAT | >100 | **<0.001** | - | - | - | - | - | - | - | - |
| SP | >100 | **<0.001** | - | - | - | - | - | - | - | - |
| SST | >100 | **<0.001** | - | - | - | - | - | - | - | - |
| SPAR | >100 | **<0.001** | - | - | - | - | - | - | - | - |
| SWTD | >100 | **<0.001** | | | | | | | | |

**Table S6. Results of posthoc test (after ANOVA) testing the impact of location and depth on different environmental parameters including metabolites, organic matter quality, nutrients and climatic conditions.** Cou = Counozouls, Man = Männikjärve, Siik = Siikaneva. Abi = Abisko, DOC = dissolved organic carbon, TN = total nitrogen, WAT = winter air temperature, WST = winter soil temperature, WP = winter precipitation, WWTD = winter water table depth, WPAR = winter PAR, SAT = spring air temperature, SST = spring soil temperature, SP = spring precipitation, SWTD = spring water table depth, SPAR = spring PAR, Cou = Counozouls, Män = Männikjärve, Siik = Siikaneva. Abi = Abisko, D1 = 0-5 cm, D2 = 5-10 cm, D3 = 10-15 cm. Bold results mean significant *p-value*.

| | Location | | | | | | Depth Cou | | | Depth Män | | | Depth Siik | | | Depth Abi | | |
|---|---|---|---|---|---|---|---|---|---|---|---|---|---|---|---|---|---|---|
| | Cou - Abi | Cou - Män | Cou - Siik | Män - Siik | Män - Abi | Abi-Siik | **D1-D2** | **D1-D3** | **D2-D3** | **D1-D2** | **D1-D3** | **D2-D3** | **D1-D2** | **D1-D3** | **D2-D3** | **D1-D2** | **D1-D3** | **D2-D3** |
| DOCq | **0.017** | 0.232 | 0.145 | 0.995 | 0.663 | 0.807 | - | - | - | - | - | - | - | - | - | - | - | - |
| Peak_A | **<0.001** | 0.846 | 0.847 | 0.368 | **<0.001** | **<0.001** | - | - | - | - | - | - | - | - | - | - | - | - |
| Peak_C | **<0.001** | 0.544 | 0.666 | **0.007** | **<0.001** | **<0.001** | - | - | - | - | - | - | - | - | - | - | - | - |
| Peak_M | **<0.001** | 0.946 | 0.07 | 0.221 | **<0.001** | **<0.001** | - | - | - | - | - | - | - | - | - | - | - | - |
| RFE | **<0.001** | 0.871 | **<0.001** | **<0.001** | **<0.001** | 0.5 | - | - | - | - | - | - | - | - | - | - | - | - |
| Freshness | **<0.001** | **<0.001** | **<0.001** | 0.219 | 0.652 | **0.015** | - | - | - | - | - | - | - | - | - | - | - | - |
| BIX | **<0.001** | **<0.001** | **<0.001** | **0.018** | 0.808 | **0.001** | - | - | - | - | - | - | - | - | - | - | - | - |
| FI | **<0.001** | 0.37 | **<0.001** | **0.03** | **0.0119** | 0.989 | - | - | - | - | - | - | - | - | - | - | - | - |
| WTD | **<0.001** | **<0.001** | **<0.001** | 0.983 | 0.633 | 0.409 | - | - | - | - | - | - | - | - | - | - | - | - |
| Tannins | 0.719 | 0.025 | **0.018** | 0.999 | 0.253 | 0.201 | 0.643 | 0.202 | 0.64 | **0.017** | **<0.001** | 0.166 | 0.382 | 0.101 | 0.662 | 0.195 | 0.435 | 0.836 |
| Water_Phenols | 0.949 | 0.926 | 0.999 | 0.914 | 0.658 | 0.959 | 0.853 | 0.976 | 0.942 | 0.999 | 0.175 | 0.185 | 0.086 | 0.183 | 0.89 | 0.758 | 0.975 | 0.632 |
| Carbohydrates | 0.901 | 0.26 | **0.001** | 0.154 | 0.652 | **0.009** | 0.26 | 0.277 | 0.999 | **0.05** | **0.005** | 0.417 | 0.935 | 0.629 | 0.829 | 0.321 | 0.165 | 0.895 |
| Flavonoids | 0.929 | 0.955 | 0.999 | 0.953 | 0.923 | 0.999 | 0.881 | 0.589 | 0.862 | 0.069 | **0.049** | 0.977 | 0.531 | 0.559 | 0.999 | 0.475 | 0.994 | 0.538 |
| Phenols | 0.935 | 0.718 | 0.776 | 0.197 | 0.966 | 0.419 | 0.5 | 0.197 | 0.773 | **0.027** | **0.007** | 0.734 | 0.434 | 0.435 | 0.999 | 0.862 | 0.977 | 0.752 |
| $Na^+$ | 0.987 | 0.997 | 0.929 | 0.847 | 0.999 | 0.781 | 0.107 | 0.089 | 0.993 | 0.685 | 0.751 | 0.993 | 0.376 | 0.975 | 0.489 | 0.989 | 0.872 | 0.801 |

| | | | | | | | | | | | | | | | | | | |
|---|---|---|---|---|---|---|---|---|---|---|---|---|---|---|---|---|---|---|
| $NH_4^+$ | 0.999 | 0.989 | 0.204 | 0.343 | 0.247 | 0.997 | 0.999 | 0.462 | 0.462 | 0.492 | 0.813 | 0.848 | 0.999 | 0.439 | 0.459 | 0.909 | 0.675 | 0.899 |
| $K^+$ | 0.305 | 0.992 | 0.998 | 0.401 | 0.967 | 0.187 | **0.017** | **0.008** | 0.915 | **0.02** | **0.0024** | 0.47 | **<0.001** | **<0.001** | 0.83 | 0.078 | **0.019** | 0.707 |
| $Mg^{2+}$ | 0.982 | **<0.001** | **0.005** | 0.243 | **<0.001** | **0.002** | 0.654 | 0.998 | 0.689 | 0.359 | 0.762 | 0.759 | 0.999 | 0.966 | 0.962 | 0.774 | 0.729 | 0.344 |
| $Ca^{2+}$ | 0.967 | 0.783 | 0.349 | 0.88 | 0.512 | 0.157 | 0.277 | 0.712 | 0.7 | 0.288 | 0.253 | 0.995 | 0.992 | 0.534 | 0.608 | 0.829 | 0.84 | 0.501 |
| $F^-$ | 0.974 | 0.432 | 0.172 | 0.944 | 0.221 | 0.071 | **0.016** | **0.044** | 0.849 | 0.947 | 0.823 | 0.642 | 0.477 | 0.994 | 0.535 | 0.424 | 0.978 | 0.537 |
| $CL^-$ | 0.764 | 0.711 | 0.944 | 0.373 | 0.999 | 0.423 | 0.517 | 0.943 | 0.71 | 0.692 | 0.744 | 0.996 | 0.999 | 0.709 | 0.692 | 0.56 | 0.652 | 0.987 |
| $NO_2^-$ | 0.542 | 0.92 | 0.999 | 0.906 | 0.894 | 0.517 | 0.19 | 0.222 | 0.994 | 0.873 | 0.607 | 0.884 | 0.999 | 0.942 | 0.939 | 0.651 | 0.994 | 0.714 |
| $NO_3^-$ | **0.066** | 0.972 | 0.999 | 0.99 | 0.167 | 0.09 | 0.268 | 0.951 | 0.403 | 0.967 | 0.054 | **0.035** | 0.939 | 0.244 | 0.389 | 0.75 | 0.878 | 0.968 |
| $BR^-$ | **0.02** | 0.463 | 0.999 | 0.463 | 0.406 | **0.02** | 0.145 | 0.036 | 0.7 | 0.603 | 0.08 | 0.371 | 0.873 | 0.067 | 0.155 | 0.969 | **0.004** | **0.003** |
| $SO_4^{2-}$ | 0.301 | 0.621 | 0.994 | 0.772 | 0.947 | 0.438 | 0.984 | 0.848 | 0.924 | 0.834 | 0.989 | 0.76 | 0.521 | 0.31 | 0.909 | 0.918 | 0.991 | 0.861 |
| $PO_4^{3-}$ | **<0.001** | **<0.001** | **0.001** | 0.971 | **<0.001** | **<0.001** | 0.308 | 0.506 | 0.918 | 0.46 | 0.348 | 0.973 | 0.991 | 0.517 | 0.591 | 0.347 | 0.266 | 0.98 |
| PH | **<0.001** | **<0.001** | **<0.001** | 0.909 | 0.068 | 0.262 | - | - | - | - | - | - | - | - | - | - | - | - |
| DOC | **<0.001** | **<0.001** | **<0.001** | 0.11 | 0.419 | **0.002** | - | - | - | - | - | - | - | - | - | - | - | - |
| TN | 0.057 | 0.005 | **<0.001** | **<0.001** | 0.798 | **<0.001** | - | - | - | - | - | - | - | - | - | - | - | - |
| WAT | **<0.001** | **<0.001** | **<0.001** | **<0.001** | **<0.001** | **<0.001** | - | - | - | - | - | - | - | - | - | - | - | - |
| WP | **<0.001** | **<0.001** | **<0.001** | **<0.001** | **<0.001** | **<0.001** | - | - | - | - | - | - | - | - | - | - | - | - |
| WST | **<0.001** | **<0.001** | **<0.001** | **<0.001** | **<0.001** | **<0.001** | - | - | - | - | - | - | - | - | - | - | - | - |
| WPAR | **<0.001** | **<0.001** | **<0.001** | **<0.001** | **<0.001** | 0.309 | - | - | - | - | - | - | - | - | - | - | - | - |
| WWTD | **<0.001** | **<0.001** | **<0.001** | **<0.001** | **<0.001** | **<0.001** | - | - | - | - | - | - | - | - | - | - | - | - |
| SAT | **<0.001** | **<0.001** | **<0.001** | **<0.001** | **<0.001** | **<0.001** | - | - | - | - | - | - | - | - | - | - | - | - |
| SP | **<0.001** | **<0.001** | **<0.001** | **<0.001** | **<0.001** | **<0.001** | - | - | - | - | - | - | - | - | - | - | - | - |
| SST | **<0.001** | **<0.001** | **<0.001** | **<0.001** | **<0.001** | **<0.001** | - | - | - | - | - | - | - | - | - | - | - | - |
| SPAR | **<0.001** | **<0.001** | **<0.001** | **<0.001** | **<0.001** | **<0.001** | - | - | - | - | - | - | - | - | - | - | - | - |
| SWTD | **<0.001** | **<0.001** | **<0.001** | **<0.001** | **<0.001** | **<0.001** | - | - | - | - | - | - | - | - | - | - | - | - |

**Table S7. Results of ANOVA testing the impact of location and depth on gene abundance.** Cou = Counozouls, Männ = Männikjärve, Siik = Siikaneva. Abi = Abisko. Bold results mean significant *p-value*.

| | Location | | Depth Cou | | Depth Män | | Depth Siik | | Depth Abi | |
|---|---|---|---|---|---|---|---|---|---|---|
| | *F-value* | *p-value* | *F-value* | *p-value* | *F-value* | *p-value* | *F-value* | *p-value* | *F-value* | *p-value* |
| 16S rRNA | 11.93 | **<0.001** | 22.16 | **<0.0001** | 2.104 | 0.165 | 0.231 | 0.798 | 0.796 | 0.473 |
| 23S rRNA | 3.489 | **0.022** | 4.109 | **0.044** | 59.45 | **<0.001** | 7.743 | **0.007** | 14.83 | **<0.001** |
| *cbbL* | 9.036 | **<0.001** | 11.13 | **0.002** | 20.52 | **<0.001** | 4.178 | **0.042** | 0.78 | 0.48 |
| *pufM* | 4.973 | **0.004** | 3.218 | **0.076** | 26.57 | **<0.001** | 2.714 | 0.107 | 5.601 | **0.019** |
| 16S rRNA (cyanobacteria) | 4.629 | **<0.001** | 4.91 | **0.028** | 58.44 | **<0.001** | 6.6 | **0.012** | 20.9 | **<0.001** |

**Table S8. Results of posthoc test (after ANOVA) testing the impact of location and depth on gene abundance.** Cou = Counozouls, Männ = Männikjärve, Siik = Siikaneva. Abi = Abisko, D1 = 0-5 cm, D2 = 5-10 cm, D3 = 10-15 cm. Bold results mean significant *p-value*.

| | Location | | | | | | Depth Cou | | | Depth Män | | | Depth Siik | | | Depth Abi | | |
|---|---|---|---|---|---|---|---|---|---|---|---|---|---|---|---|---|---|---|
| | Cou - Män | Cou - Siik | Cou - Abi | Män - Siik | Män - Abi | Abi - Siik | **D1 - D2** | **D1 - D3** | **D2 - D3** | **D1 - D2** | **D1 - D3** | **D2 - D3** | **D1 - D2** | **D1 - D3** | **D2 - D3** | **D1 - D2** | **D1 - D3** | **D2 - D3** |
| 16S RNA | **<0.001** | **<0.001** | **0.016** | 0.985 | 0.122 | 0.239 | **0.029** | **<0.001** | **0.008** | 0.942 | 0.176 | 0.287 | 0.906 | 0.967 | 0.784 | 0.442 | 0.833 | 0.779 |
| 23S RNA | **0.015** | 0.349 | 0.105 | 0.483 | 0.861 | 0.915 | **0.04** | 0.624 | 0.189 | **<0.001** | **<0.001** | **<0.001** | 0.588 | **0.007** | **0.04** | **0.007** | **<0.001** | 0.314 |
| *cbbL* | **<0.001** | **0.002** | 0.119 | 0.674 | **0.049** | 0.432 | **0.042** | **0.001** | 0.175 | 0.789 | **<0.001** | **<0.001** | 0.14 | 0.756 | **0.04** | 0.556 | 0.531 | 0.999 |
| *pufM* | **0.003** | **0.05** | 0.069 | 0.699 | 0.621 | 0.999 | 0.415 | 0.063 | 0.463 | **0.048** | **<0.001** | **0.001** | 0.945 | 0.199 | 0.119 | 0.204 | 0.055 | 0.316 |
| 16S rRNA (cyanobacteria) | **0.006** | 0.23 | **0.022** | 0.45 | 0.97 | 0.73 | **0.027** | 0.71 | 0.11 | **<0.001** | **<0.001** | **<0.001** | 0.78 | **0.013** | **0.043** | **<0.001** | **<0.001** | 0.99 |

**Table S9. Results of ANOVA testing the impact of location and depth on 23S rRNA gene, *cbbL* gene and *bchY* gene alpha richness (Chao1 index) and alpha diversity (Shannon index).** Cou = Counozouls, Männ = Männikjärve, Siik = Siikaneva. Abi = Abisko. Bold results mean significant *p-value*.

| | | Location | | Depth Cou | | Depth Män | | Depth Siik | | Depth Abi | |
|---|---|---|---|---|---|---|---|---|---|---|---|
| | | *F-value* | *p-value* | *F-value* | *p-value* | *F-value* | *p-value* | *F-value* | *p-value* | *F-value* | *p-value* |
| 23S rRNA gene (oxygenic phototrophs) | Chao1 | 7.537 | **<0.001** | 7.676 | **0.007** | 23.59 | **<0.001** | 0.44 | 0.652 | 0.913 | 0.427 |
| | Shannon | 1.817 | 0.155 | 6.804 | **0.012** | 15.83 | **<0.001** | 1.475 | 0.267 | 1.035 | 0.385 |
| *cbbL* gene (chemoautotrophs) | Chao1 | 14.58 | **<0.001** | 1.94 | 0.186 | 104.7 | **<0.001** | 5.941 | **0.016** | 18.84 | **<0.001** |
| | Shannon | 4.07 | **0.011** | 7.812 | **0.007** | 66.79 | **<0.001** | 2.963 | 0.09 | 15.07 | **<0.001** |
| *bchY* gene (AAnPBs) | Chao1 | 3.413 | **0.024** | 1.969 | 0.182 | 16.02 | **<0.001** | 6.196 | **0.014** | 1.695 | 0.225 |
| | Shannon | 9.767 | **<0.001** | 2.389 | 0.134 | 5.552 | **0.02** | 15.55 | **<0.001** | 2.554 | 0.119 |

**Table S10. Results of posthoc test (after ANOVA) testing the impact of location and depth on 23S rRNA gene, *cbbL* gene and *bchY* gene richness (chao1 index) and alpha diversity (Shannon index).** Cou = Counozouls, Männ = Männikjärve, Siik = Siikaneva. Abi = Abisko, D1 = 0-5 cm, D2 = 5-10 cm, D3 = 10-15 cm. Bold results mean significant *p-value*.

| | | Location | | | | | | Depth Cou | | | Depth Män | | | Depth Siik | | | Depth Abi | | |
|---|---|---|---|---|---|---|---|---|---|---|---|---|---|---|---|---|---|---|---|
| | | Cou - Män | Cou - Siik | Cou - Abi | Män - Siik | Män - Abi | Abi - Siik | **D1 - D2** | **D1 - D3** | **D2 - D3** | **D1 - D2** | **D1 - D3** | **D2 - D3** | **D1 - D2** | **D1 - D3** | **D2 - D3** | **D1 - D2** | **D1 - D3** | **D2 - D3** |
| 23S rRNA gene (oxygenic phototrophs) | Chao1 | **<0.001** | 0.107 | **0.002** | 0.196 | 0.958 | 0.439 | **0.009** | **0.025** | 0.824 | 0.998 | **<0.001** | **<0.001** | 0.941 | 0.823 | 0.633 | 0.901 | 0.657 | 0.409 |
| | Shannon | 0.827 | 0.613 | 0.799 | 0.982 | 0.3 | 0.154 | **0.021** | **0.018** | 0.996 | 0.671 | **0.002** | **<0.001** | 0.838 | 0.249 | 0.522 | 0.78 | 0.353 | 0.734 |
| *cbbL* gene (chemoautotrophs) | Chao1 | 0.322 | **<0.001** | 0.124 | **<0.001** | 0.953 | **<0.001** | 0.924 | 0.191 | 0.332 | **0.032** | **<0.01** | **<0.01** | 0.814 | 0.053 | **0.018** | 0.317 | **0.002** | **0.002** |
| | Shannon | 0.966 | **0.013** | 0.266 | **0.045** | 0.521 | 0.557 | 0.26 | **0.005** | 0.098 | **0.008** | **<0.001** | **<0.001** | 0.763 | 0.084 | 0.259 | 0.725 | **<0.001** | **0.003** |
| *bchY* gene (AAnPBs) | Chao1 | 0.149 | 0.987 | 0.855 | 0.073 | **0.023** | 0.966 | 0.954 | 0.197 | 0.302 | 0.6 | **0.04** | **0.04** | **0.049** | 0.524 | **0.012** | 0.04 | 0.198 | 0.637 |
| | Shannon | **<0.001** | **0.006** | 0.937 | 0.689 | **0.0011** | **0.028** | 0.447 | 0.116 | 0.634 | 0.061 | 0.788 | 0.057 | **<0.001** | **0.0023** | 0.72 | 0.791 | 0.111 | 0.307 |

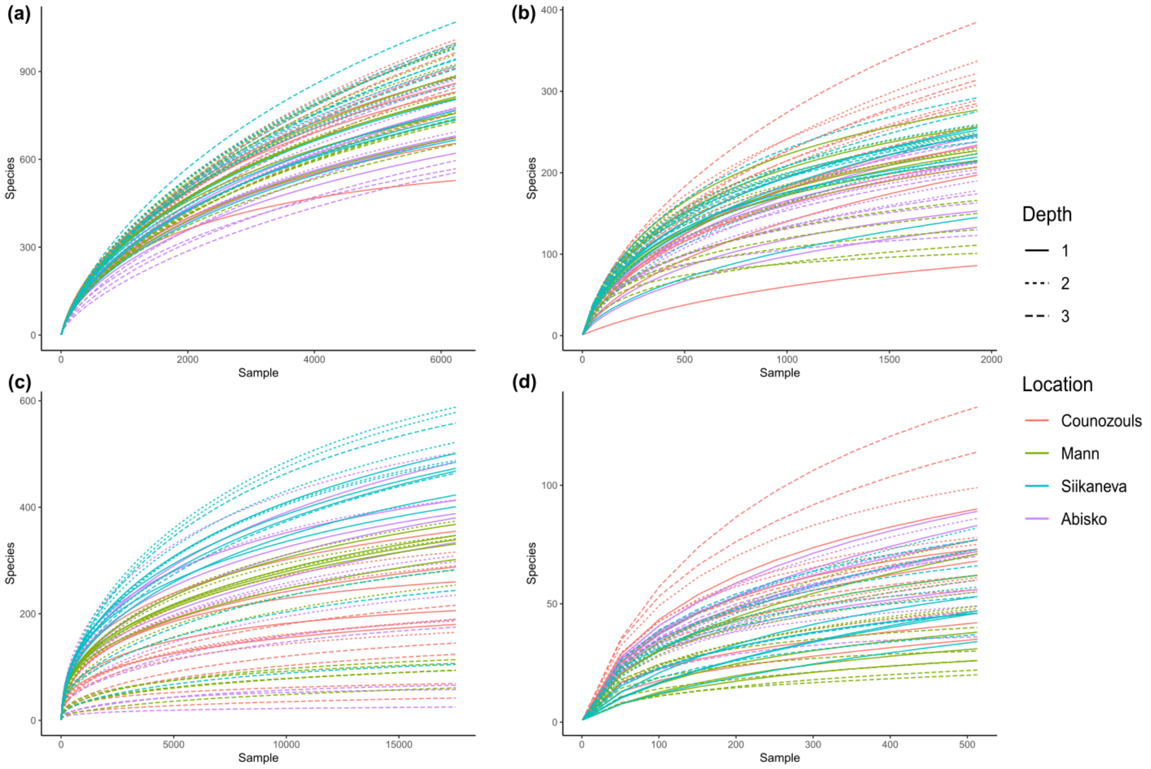


**Figure S1. Rarefaction curves** of microbial sequences after normalization of the number of sequences for **(a)** 16S rRNA gene, **(b)** 23S rRNA gene, **(c)** *cbbL* gene and **(d)** *bchY* gene.

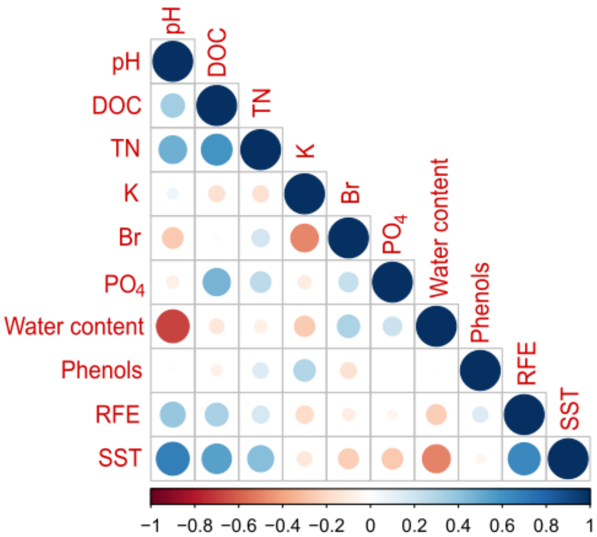


**Figure S2. Correlation plot of environmental variables** retained for ACP analysis. DOC = dissolved organic carbon, TN = total nitrogen, RFE = relative fluorescence efficiency and SST = spring soil temperature.

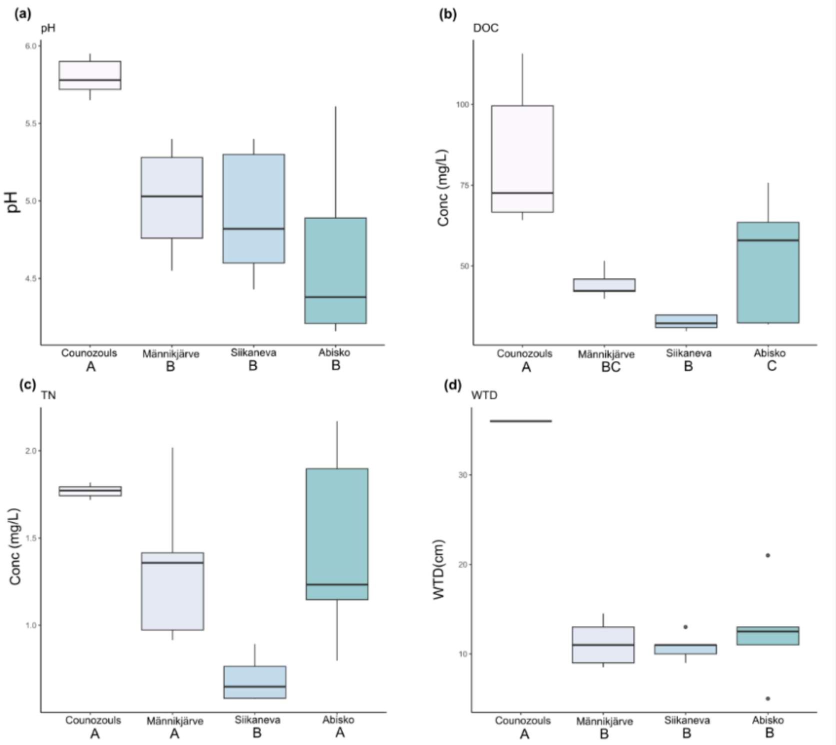


**Figure S3. Boxplot describing (a) pH, (b) DOC, (c) TN and (d) WTD for each location.** DOC = dissolved organic carbon, TN = total nitrogen and WTD = water table depth. Uppercase letters represent the differences between each peatland.

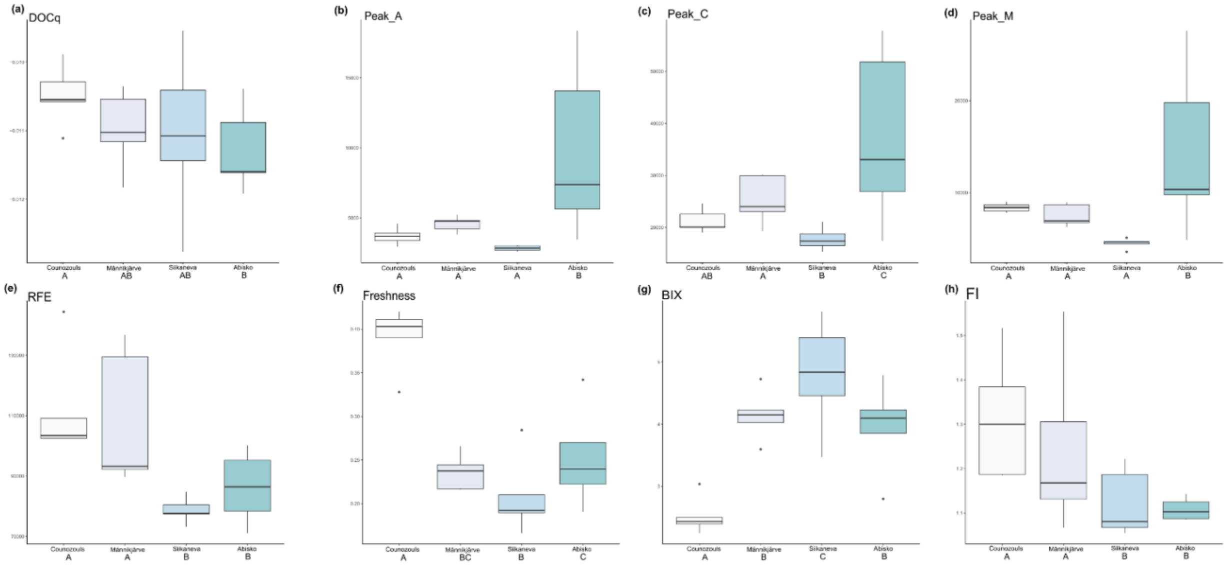


**Figure S4. Boxplot describing organic matter quality index (DOCq, peak_A, peak_C, peak_M, RFE, freshness, BIX and FI) for each location.** Uppercase letters represent the differences between each peatland. DOC = dissolved organic carbon, RFE = relative fluorescence efficiency, BIX = biological index and FI = fluorescence index.

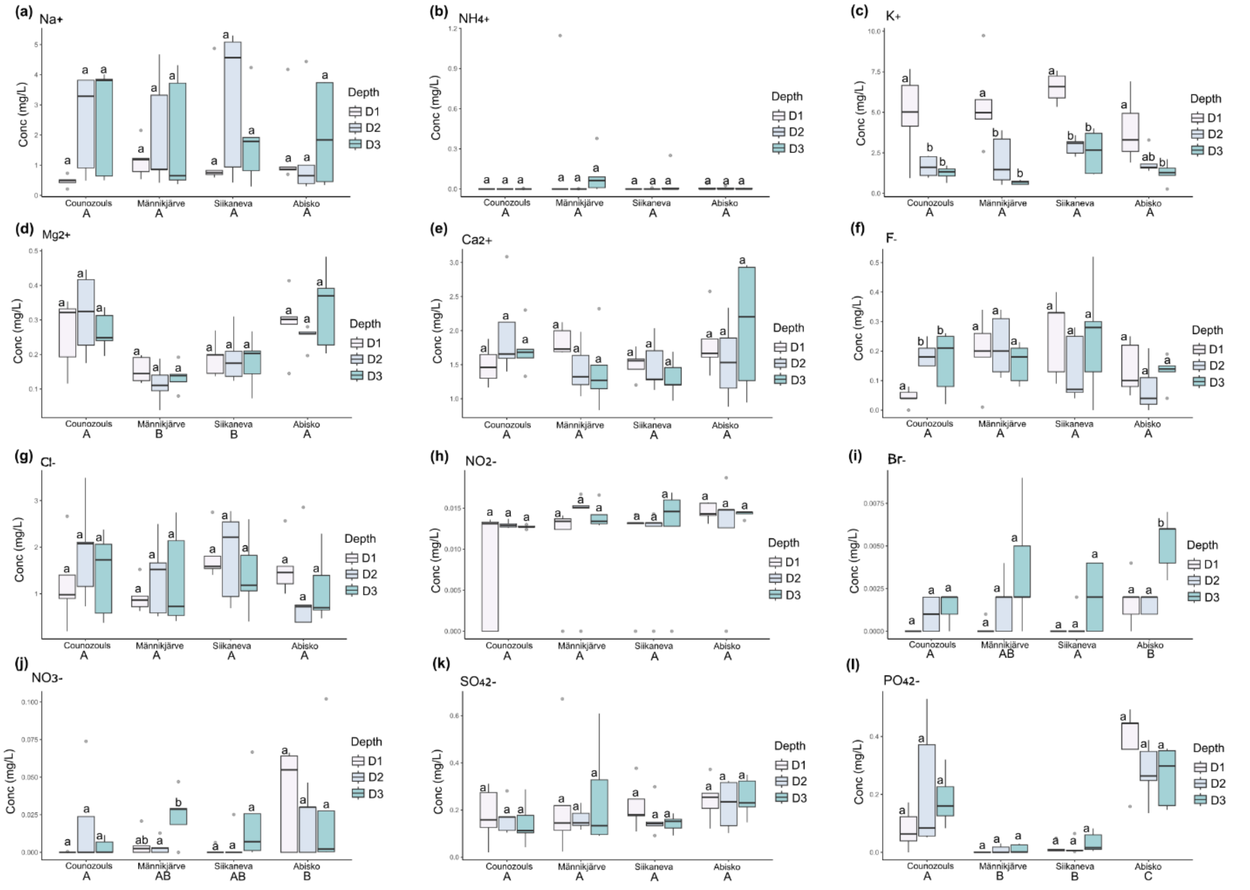


**Figure S5. Boxplot describing cations (**$Na^{+}$**,** $NH_4^{+}$**,** $K^{+}$**,** $Mg^{2+}$ **and** $Ca^{2+}$**) and anions (**$F^{-}$**,** $Cl^{-}$**,** $NO_2$**,** $NO_3^{-}$**,** $Br^{-}$**,** $SO_4^{2-}$ **and** $PO_4^{3-}$**) for each location.** Uppercase letters represent the differences between each peatland, lowercase letters represent the differences between depth (D1, D2 and D3) at each location. D1 = 0-5 cm, D2 = 5-10 cm and D3 = 10-15 cm.

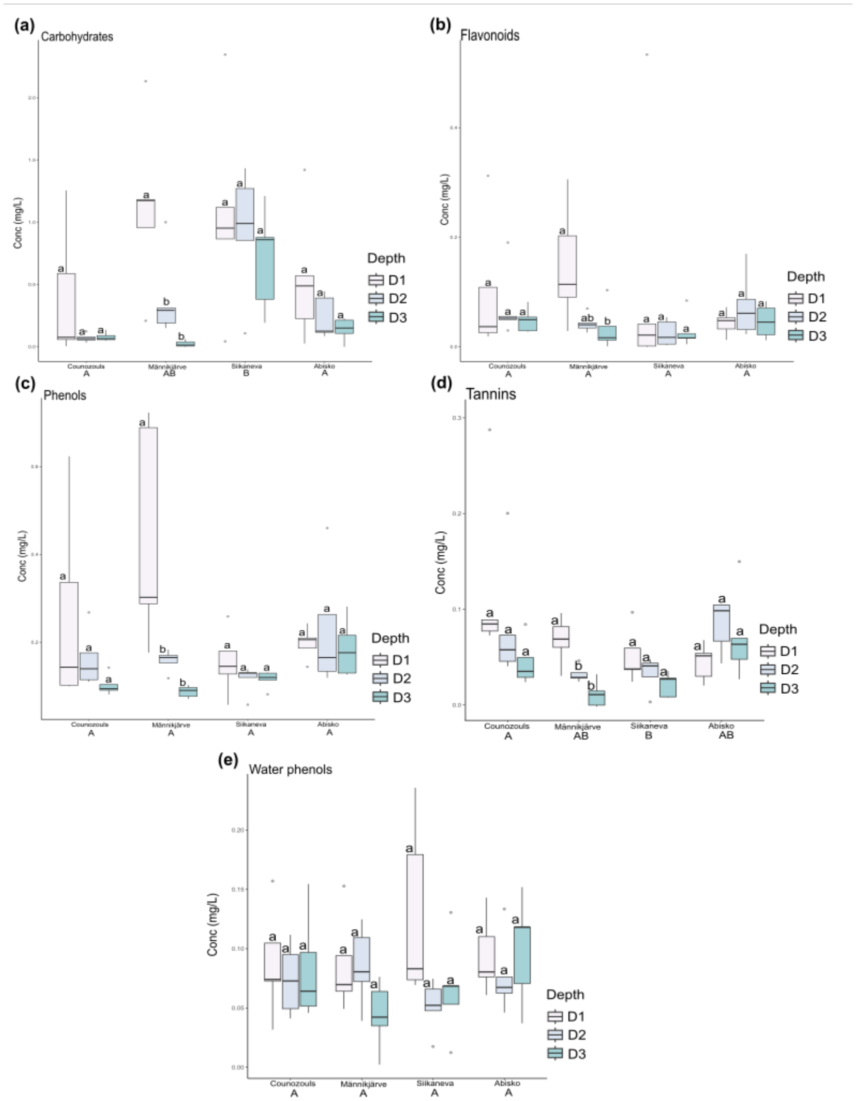


**Figure S6. Boxplot describing metabolites (carbohydrates, flavonoids, phenols, tannins and water phenols) for each location.** Uppercase letters represent the differences between each peatland, lowercase letters represent the differences between depth (D1, D2 and D3) at each location. D1 = 0-5 cm, D2 = 5-10 cm and D3 = 10-15 cm.

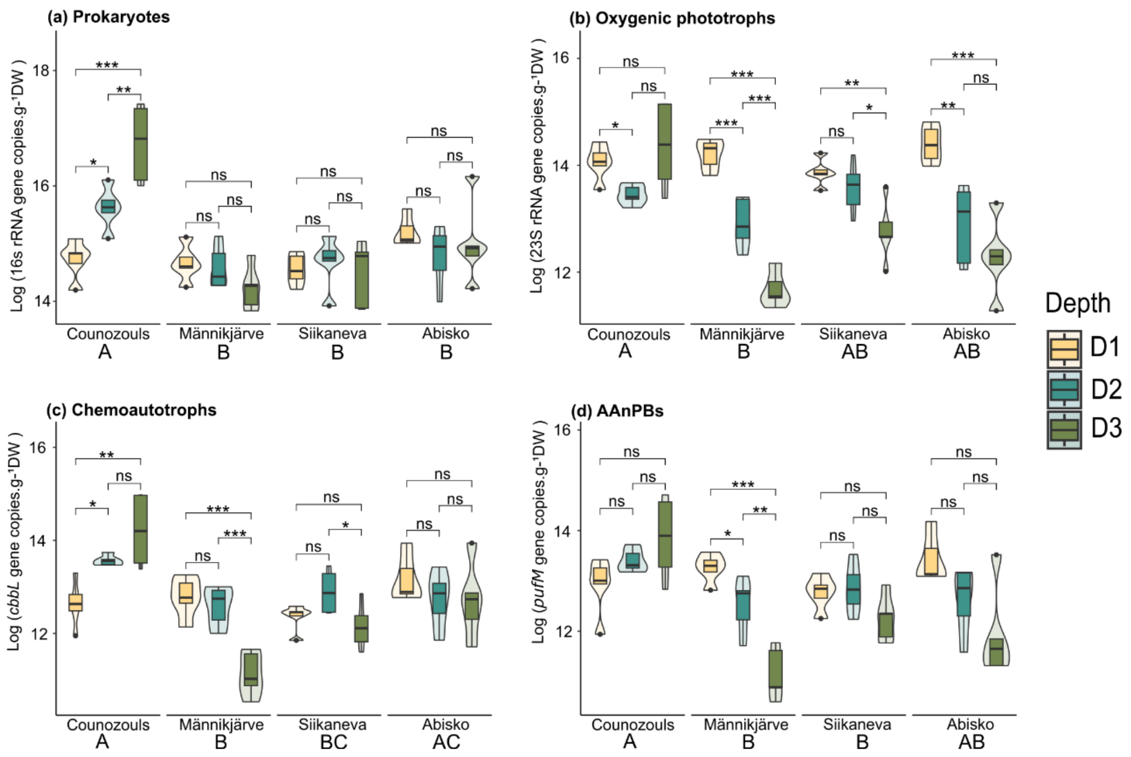


**Figure S7. Absolute quantification using ddPCR** of 16S rRNA gene (**(a)**; prokaryotes), 23S rRNA gene (**(b)**; oxygenic phototrophs), *cbbL* (**(c)**; (chemoautotrophs) and *pufM* (**(d)**; AAnPBs) at different depths in the four peatlands. Violin plots are showing the data distribution shape while boxplots are representing the logarithm of the total gene copies.g$^{-1}$ DW. D1 = 0-5 cm; D2 = 5-10 cm and D3 = 10-15 cm. ns = not significant, * = 0.05 P < 0.01 , ** = 0.01 < P < 0.001 and *** = P < 0.001.

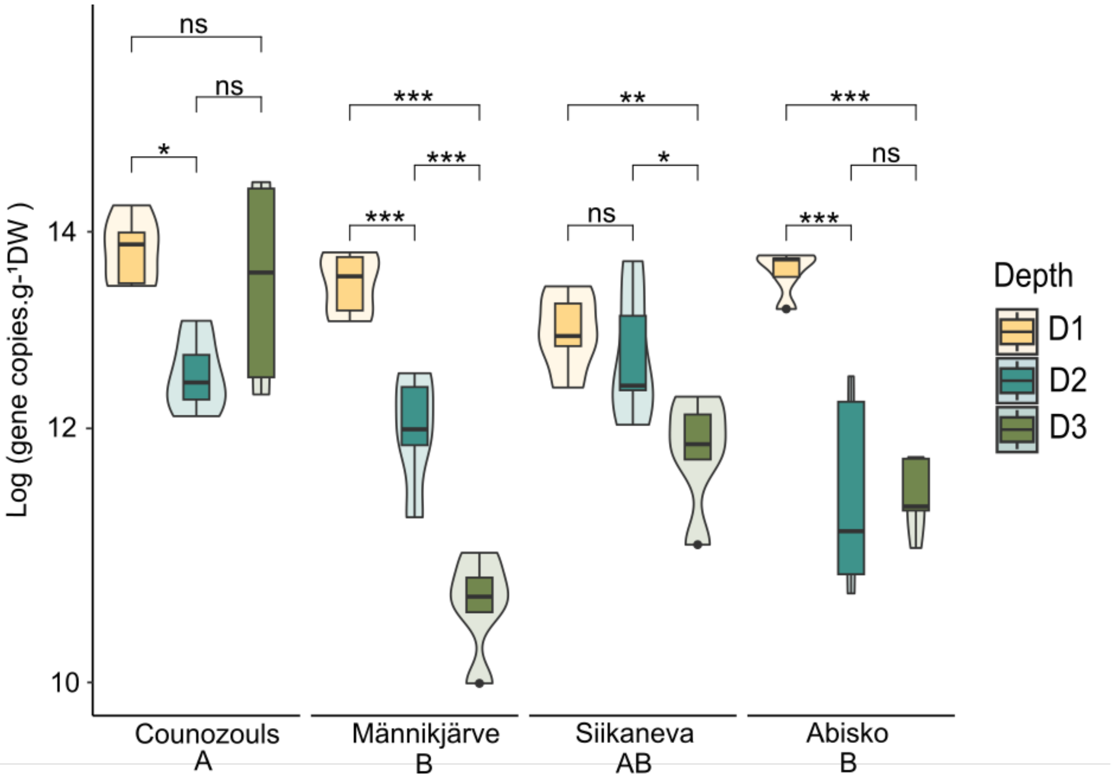


**Figure S8. Absolute quantification of cyanobacteria using ddPCR.** Violin plots are showing the data distribution shape while boxplots are representing the logarithm of the total gene copies.g$^{-1}$ DW. D1 = 0-5 cm; D2 = 5-10 cm and D3 = 10-15 cm. ns = not significant, * = 0.05 P < 0.01 , ** = 0.01 < P < 0.001 and *** = P < 0.001.

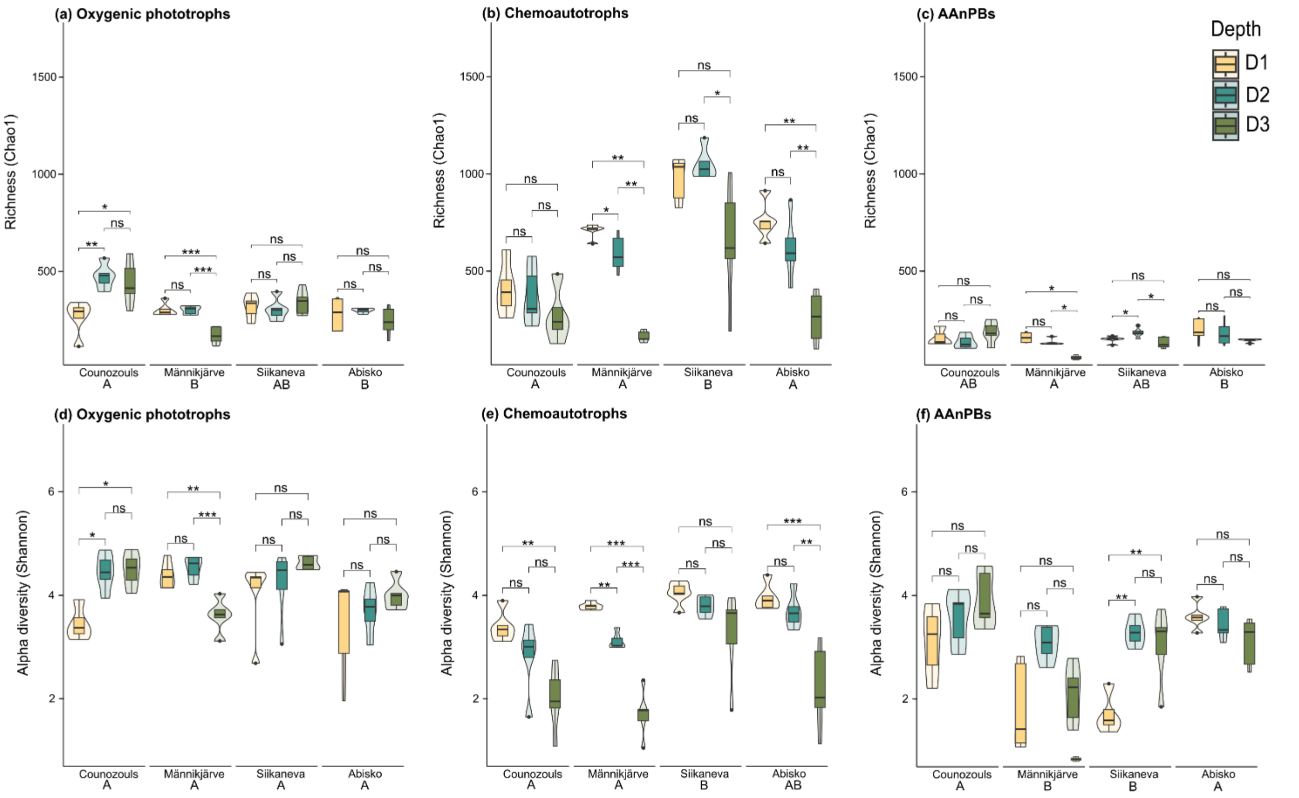


**Figure S9. Diversity description** of 23S rRNA, *cbbL* and *bchY* genes at each location and depth (D1, D2 and D3). **(a), (b) and (c)** Richness (Chao1 index) and **(d), (e) and (f)** Alpha diversity (Shannon index). Violin plots are showing the data distribution shape while boxplots are representing the logarithm of the total gene copies.g$^{-1}$ DW. D1 = 0-5 cm, D2 = 5-10 cm and D3 = 10-15 cm. ns = not significant, * = 0.05 P < 0.01 , ** = 0.01 < P < 0.001 and *** = P < 0.001.

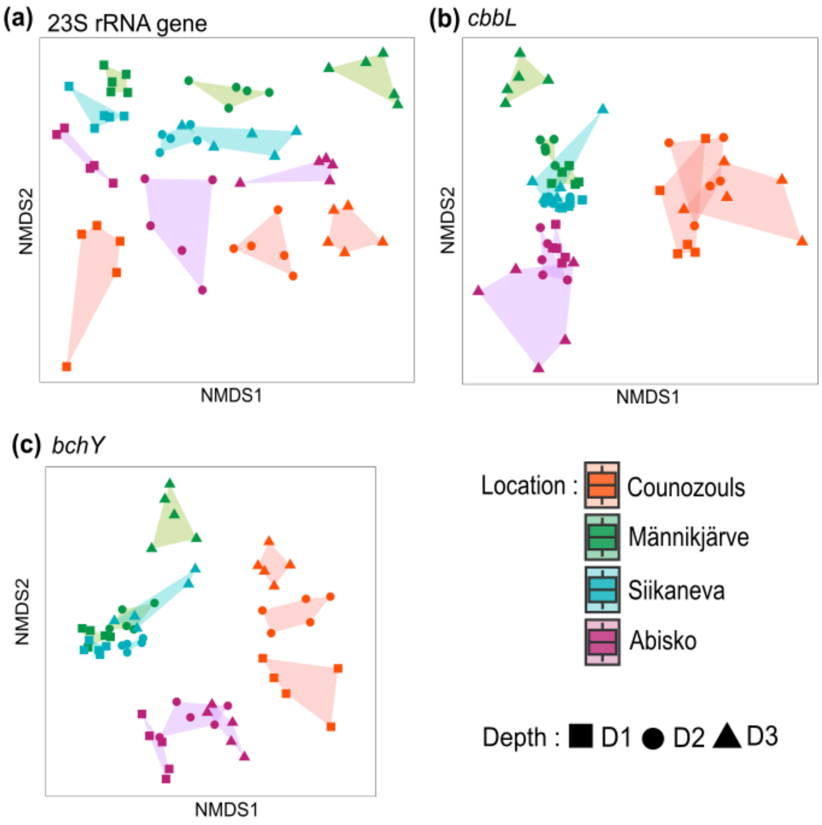


**Figure S10. NMDS showing the community structure** of **(a)** oxygenic phototrophs (23S rRNA gene), **(b)** chemoautotrophs (cbbL gene) and **(c)** AAnPBs (bchY gene). The Bray-Curtis dissimilatory index has been used. D1 = 0-5 cm; D2 = 5-10 cm and D3 = 10-15 cm.

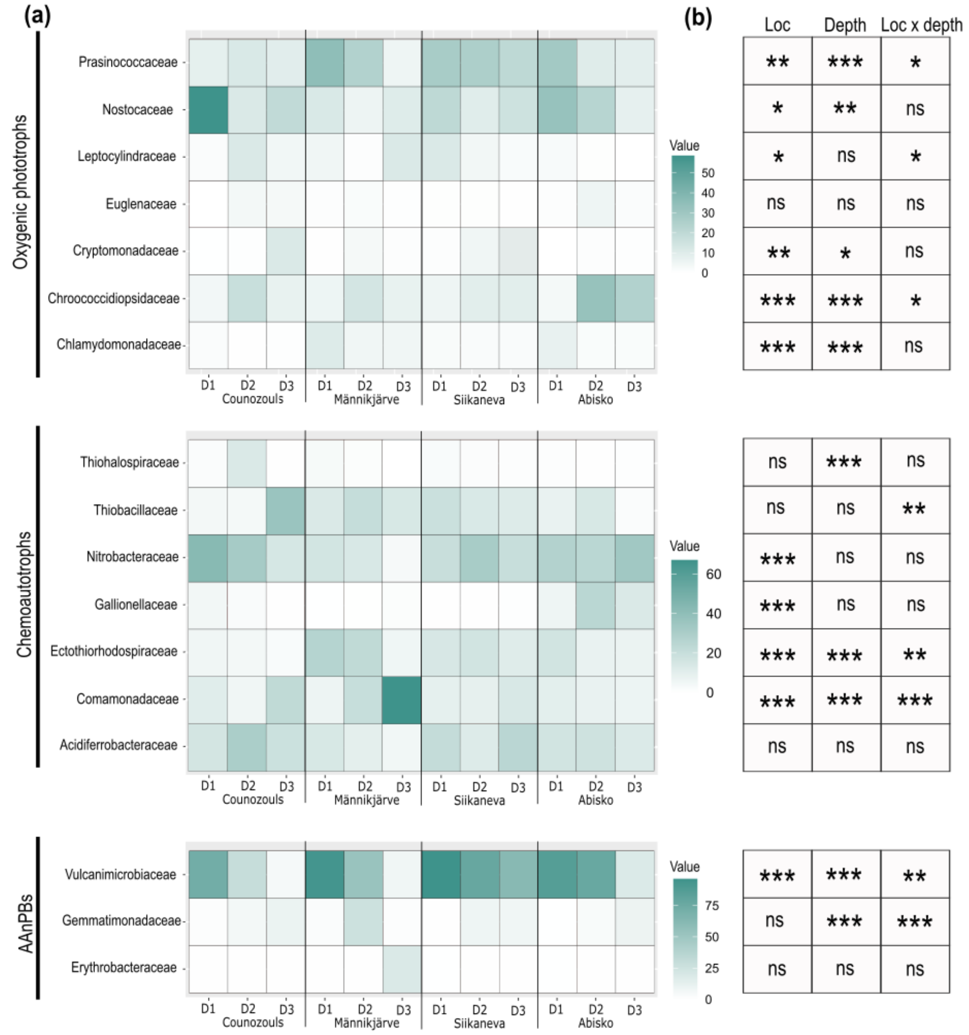


**Figure S11. Impact of location and depth on relative abundance of ASVs aggregated by family. (a)** Heatmaps showing the relative abundance of ASVs aggregated by family according to location and depth. Only family with abundance higher than 5% were kept. Light red represents low abundances while dark red represents higher abundances. D1 = 0-5 cm, D2 = 5-10 cm and D3 = 10-15 cm. **(b)** P-values of the explanatory power of location, depth and location with depth. Loc = location, * = 0.01 < *P* < 0.05; ** = 0.001 < *P* < 0.01 and *** = *P* < 0.01.

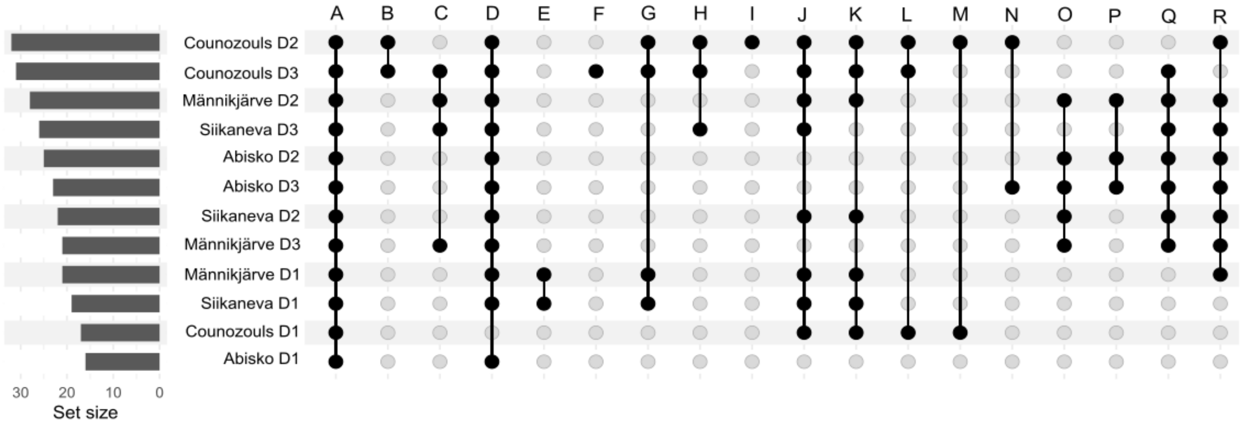


| | |
|---|---|
| A | Bacillariophyceae, Synurophyceae, Trebouxiophyceae, Euglenophyceae, Mediophyceae, Palmophyllophyceae, Cryptophyceae, Cyanophyceae, |
| B | Anthocerotopsida, Myxococcia, Omnitrophia |
| C | Porphyridiophyceae |
| D | Firestonebacteria, Defferrisomatia |
| E | Glaucophyceae |
| F | Actinobacteria |
| G | Dinophyceae |
| H | Bacteroidia |
| I | Clostridia, Leiosporocerotopsida |
| J | Ulvophyceae |
| K | Conjugatophyceae |
| L | Xanthophyceae |
| M | Synergistia |
| N | Clostridia |
| O | Planctomycetes |
| P | Florideophyceae |
| Q | Porphyridiophyceae |
| R | Desulfuromonodales |

**Figure S12. Upset plot of the presence of ASVs aggregated by class for the 23S rRNA gene in the four peatland sites.** D1 = 0-5 cm, D2 = 5-10 cm and D3 = 10-15 cm.

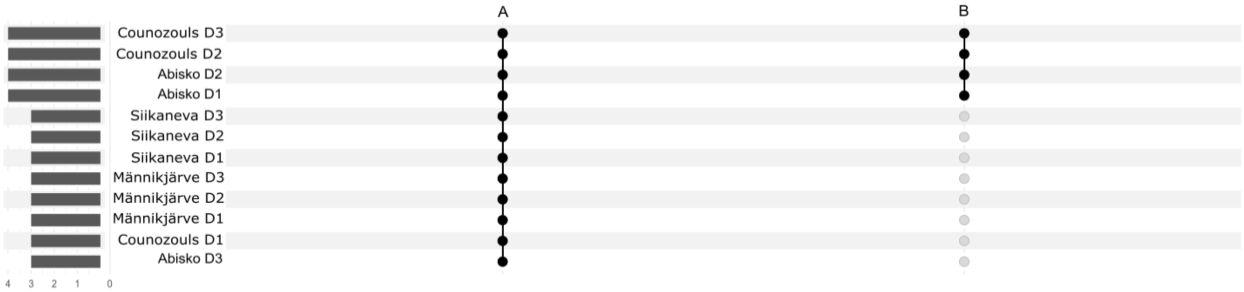


| | |
|---|---|
| A | Alphaproteobacteria, Betaproteobacteria, Gammaproteobacteria |
| B | unassigned |

**Figure S13. Upset plot of the presence of ASVs aggregated by class for the *cbbL* gene in the four peatland sites.** D1 = 0-5 cm, D2 = 5-10 cm and D3 = 10-15 cm.

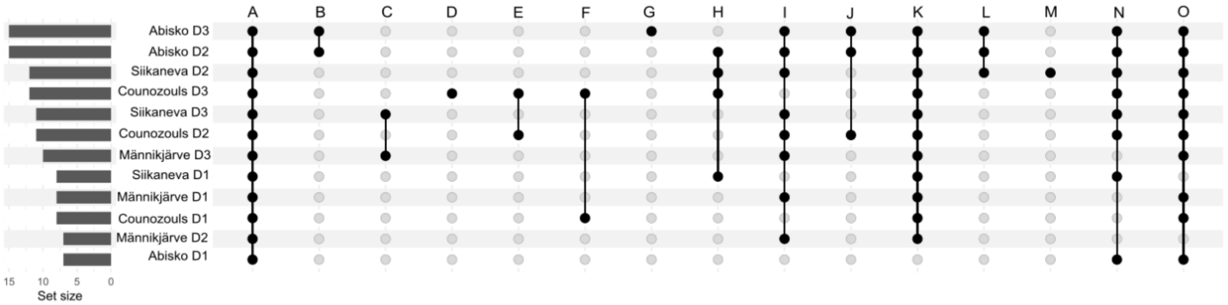


| | |
|---|---|
| A | Vulcanimicrobiia, unassigned, Gemmatimonadetes, Terriglobia, Alphaproteobacteria |
| B | Chitinophagia, Thermoleophilia, Vicinamibacteria |
| C | Deinococcota, Desulfovibrionia |
| D | Longimicrobiia |
| E | Deltaproteobacteria |
| F | Chloroflexia |
| G | Desulfobacteria |
| H | Desulfuromonadia |
| I | Actinomycetota |
| J | Acidimicrobiia |
| K | Betaproteobacteria |
| L | Actinomycetes |
| M | Myxococcia |
| N | Gammaproteobacteria |
| O | Planctomycetia |

**Figure S14. Upset plot of the presence of ASVs aggregated by class for the *bchY* gene in the four peatland sites.** D1 = 0-5 cm, D2 = 5-10 cm and D3 = 10-15 cm.

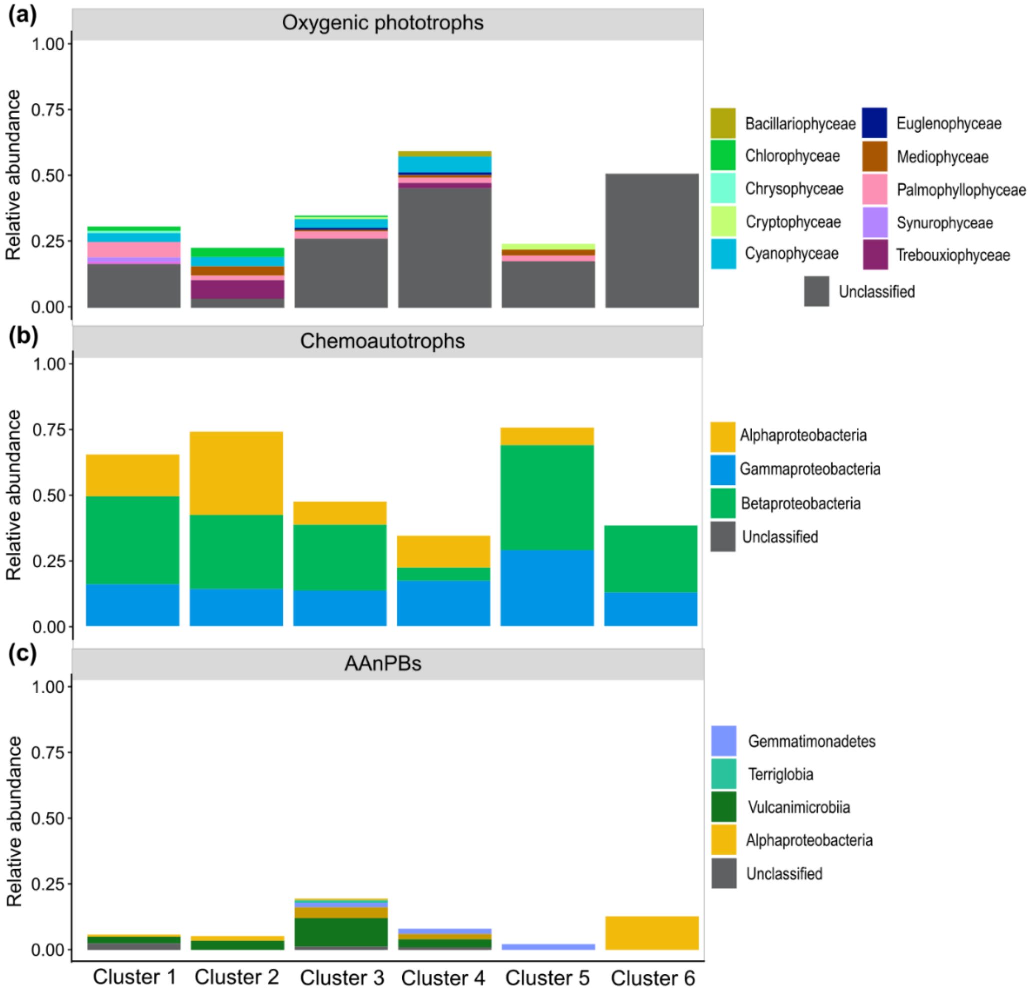


**Figure S15. Barplot of the relative abundance of each class constituting the different clusters** for **(a)** oxygenic phototrophs, **(b)** chemoautotrophs and **(c)** AAnPBs.

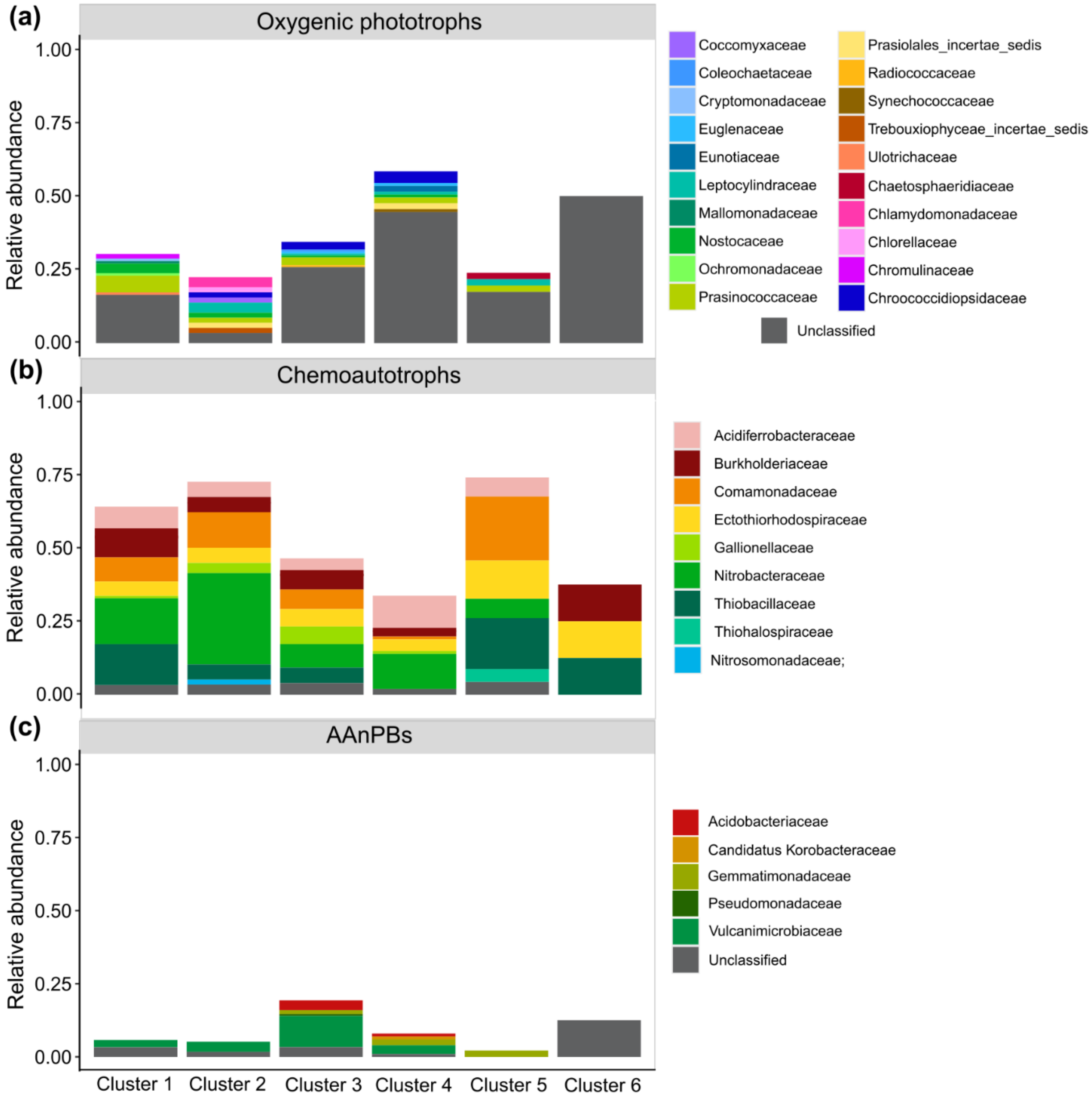


**Figure S16. Barplot of the relative abundance of each family constituting the different clusters** for **(a)** oxygenic phototrophs, **(b)** chemoautotrophs and **(c)** AAnPBs.
